# Auditory cortical coding of spectrotemporal modulations is sufficient to support perceptual categorization of speech and music

**DOI:** 10.1101/2025.06.05.657930

**Authors:** Jérémie Ginzburg, Clarisse Gotti, Émilie Cloutier Debaque, Arthur Borderie, Benjamin Morillon, Laurence Martineau, Paule Lessard Bonaventure, Robert J Zatorre, Philippe Albouy

## Abstract

Auditory processing is typically described as hierarchical, culminating in neural representation of abstract categories. However, it remains unclear whether category-selective responses in auditory cortex require representational mechanisms beyond the coding of acoustic features, or whether the acoustic representations already available in the auditory cortex are sufficient to account for categorization. Here, we test whether cortical coding of spectrotemporal modulation (STM) features is sufficient to support speech–music categorization by combining human intracranial recordings with continuous behavioral judgments of a naturalistic soundtrack in which speech and music occur both separately and simultaneously. We show that temporal and spectral modulation patterns largely characterize speech and music, respectively, and that cortical auditory regions robustly track these features over time, with distinct oscillatory frequency bands preferentially encoding temporal and spectral modulations. Critically, cortical representations of STMs predicted perceptual categorical judgments gathered in an independent sample. Finally, speech- and music-related STM representations showed stronger tracking of category-specific acoustical features in left versus right cortical auditory regions, respectively. These findings indicate that the efficient neural coding of acoustical features provides a sufficient basis for the categorical distinction between speech and music, and that the temporal and spectral components of this representation are implemented through distinct oscillatory mechanisms.

## INTRODUCTION

The conversion of sensory input into behaviorally relevant and stable perceptual objects is a core property of brain function (*1–3*). In audition, this conversion has been described as a hierarchical progression in which the physical sound waveforms are successively transformed into increasingly complex and progressively more abstract representations (*4–6*). The auditory cortex, in particular, is known to represent sounds as a function of their spectrotemporal modulation (STM) structure in both humans and other species (*7–16*). Speech and music are arguably the two most cognitively complex ways that humans use sound. Yet, a central open question is how much of the categorical distinction between speech and music can be accounted for by STM coding itself, and whether category-selective responses in the auditory cortex require representational mechanisms that go beyond the coding of acoustic features. These two possibilities are not mutually exclusive, since acoustic and categorical representations may coexist within a hierarchical system, but they differ in the explanatory weight that they assign to acoustic coding. Previous studies have approached this question from contrasting perspectives (*7*, *11*, *17–23*).

This question carries significant implications for our understanding of brain function, including the question of whether higher-order representations are localized or distributed across broader networks (*4*, *24*, *25*). Some neuroimaging studies have emphasized functional specialization, reporting bilateral cortical regions that respond specifically to speech, song, or instrumental music, with selectivity that was not accounted for by the STM features against which it was tested (*21*, *22*, *26*, *27*). Other work has emphasized that the distinction between speech and music can be captured by consistent differences in the acoustical cues that characterize these two classes of sounds, which are, in addition, processed with different hemispheric weighting (*7*, *17*, *28–33*). These two lines of work are thus assigning different explanatory weight to acoustic coding.

Several recent influential studies have specifically suggested the human auditory cortex is organized to support specialized processing pathways for behaviorally meaningful sound categories. According to this framework, while processing within early stages of the auditory pathway can be largely explained by tuning to basic acoustic features, category-specific representations emerge beyond these early areas, where neural processing becomes increasingly non-linear (*19*). These transformations are thought to give rise to representations in non-primary auditory regions that encode auditory objects in ways that are more invariant to low-level acoustic variability (*34*). Supporting this view, several studies have reported distinct neural populations in bilateral non-primary auditory areas that respond selectively to speech, music, or even specifically to songs (*19*, *21*, *22*, *35*, *36*), concluding that these responses were not explainable by the STM features against which they were evaluated (*21–23*).

While these findings suggest that higher-level auditory areas may host representations that go beyond the acoustic input, it remains an open question whether such categorical responses require representational mechanisms operating beyond the acoustic feature coding that is implemented within the established auditory processing hierarchy. From this perspective, categorical distinctions between sound classes may not necessitate distinct neural modules dedicated to each category but could instead be supported, at least in substantial part, by the way the auditory system organizes and transforms acoustic information.

A promising framework for addressing this question is provided by the STM account. The concept of spectrotemporal receptive fields (*5*) offers a computationally grounded and neurophysiologically plausible framework for understanding how auditory neurons decompose incoming acoustic signals (*13*, *14*, *16*). This model suggests that auditory neurons function as STM rate filters, a proposal supported by single-cell recordings in animals (*8*, *10*, *15*, *16*, *37*), and human neuroimaging studies (*11*, *12*, *38*, *39*). A recent behavioral study has also shown that STM-based representations are sufficient to capture spectrotemporal features that can robustly discriminate song from speech across diverse cultures, outperforming a wide range of other acoustic descriptors (*17*). Furthermore, the STM framework has been shown to account for the functional hemispheric lateralization of speech and music processing (*7*, *29*), and that this lateralization, rooted in the STM encoding scheme, extends beyond speech and music to other types of sounds (*28*, *30–33*). Taken together, these observations motivate the present direct evaluation of whether the STM representations observed in the auditory cortex are sufficiently structured to account for the emergence of perceptual categories such as speech and music.

The question of whether STM representations are sufficient to support categorization is closely tied to the question of how these representations are implemented in cortical activity, which remains only partly characterized. The evidence that the auditory cortex tracks STM features comes in large part from intracranial recordings in which the neural signal is summarized by the amplitude of high-frequency activity, for instance in the range of 70 to 150 Hz. In this context, high-frequency activity is used as a proxy for local population spiking (*22*, *40*) and has allowed the reconstruction of speech and music spectrograms and the estimation of spectrotemporal receptive fields directly from cortical activity (*9*, *38*, *41*). The contribution of the other frequency bands is of particular relevance for the STM framework, because the temporal and the spectral dimensions of the STM representation differ in the timescales over which the acoustic signal has to be integrated, and because cortical oscillations operating at different rates have been proposed to provide the computational units within which acoustic information is sampled and analyzed (*42*, *43*). Consistent with this view, a recent intracranial study covering the entire frequency spectrum of the neurophysiological signal has shown that low-frequency activity encodes the temporal dynamics of continuous speech and music in a larger number of recording sites than high-frequency activity does (*44*). Because the acoustic descriptors used in that work (i.e., temporal envelope and its acoustic onset edges) characterize the signal only along its temporal dimension, it remains unknown whether the spectral dimension of the acoustic signal is supported by the same oscillatory substrate. The STM decomposition provides a direct means of addressing this question, since it resolves the temporal and the spectral dimensions of the acoustic signal explicitly and therefore allows the contribution of each frequency band to be estimated for each dimension separately.

The present study aimed to determine whether the cortical representation of the STM features of naturalistic sounds contains sufficient information to account for the perceptual categorization of speech and music. We recorded stereo-electroencephalography (sEEG) activity in auditory cortex via implanted electrodes from patients with intractable epilepsy while they passively listened to a naturalistic 7-minute excerpt from the audio track of a movie. The excerpt contained speech and music presented both separately and simultaneously, allowing us to approach natural listening conditions with neural measures of high spatial and temporal precision. This approach differs from the designs on which the STM framework has so far been evaluated, in which brief, artificial speech and music excerpts are presented in isolation and a single modulation spectrum is computed for each excerpt. Under real-life continuous listening conditions, the two categories are interleaved and frequently superimposed, their boundaries are not marked, and the acoustic content changes from moment to moment, so that the STM decomposition has to be resolved in time and the question becomes whether the cortical representation supports categorization as it unfolds rather than whether two isolated classes of sounds can be told apart. Separately, behavioral categorization data were collected from healthy participants listening to the same excerpt to identify segments of speech and/or music. Using ridge regression and temporal response function models to track STM features over time, we tested whether intracranial oscillatory activity could predict the STM features of the audio track, and whether these features were in turn sufficient to predict behavioral categorization in the independent group (out-of-sample prediction). Because the neural representation used in the latter step is restricted to STM coordinates, this approach measures the amount of categorical information carried by an acoustically grounded cortical representation.

We formulated six key predictions: (i) If the behavioral categorization of the continuous naturalistic stimulus is based on STM features, we should observe distinct STM patterns in the acoustics of the waveform for segments classified as speech and music, replicating findings from studies using shorter and more artificial stimuli (*7*, *17*). (ii) If the brain tracks these STM patterns in naturalistic settings, they should be robustly encoded in the auditory cortex, particularly in non-primary regions (*7*, *45*). (iii) If the temporal and spectral dimensions of the STM representation are integrated over different timescales, their tracking should be supported by a structured contribution of distinct cortical frequency bands. (iv) If STM features alone are sufficient to support perceptual categorization, then brain-reconstructed STMs (i.e., STMs predicted from neural signals) should be able to account for behavioral categorization judgments obtained from an independent group of participants. (v) Under the same assumption, the brain-reconstructed STMs should closely match the acoustically derived STMs that are most relevant for categorization in the healthy listeners, as determined in the previous analysis step. (vi) Finally, the similarity between brain-reconstructed and acoustically derived STM patterns computed in (v) should exhibit hemispheric asymmetries, stronger in the left hemisphere for speech/temporal modulations, and in the right hemisphere for music/spectral modulations, in line with the literature.

## RESULTS

### STM features predict perceptual categorization: spectral modulations for music, temporal modulations for speech

We collected behavioral categorization data from 19 healthy participants listening to the first 7 minutes and 50 seconds of the French-dubbed audio track of *Harry Potter and the Philosopher’s Stone*. Participants continuously indicated whether they perceived speech and/or music by pressing corresponding keys (see Figure 1A and Methods), resulting in binary vectors for each participant reflecting perceived categories over time. Overlapping responses were allowed by design, as speech and music co-occur frequently in the audio track, and capturing such overlap provides a more ecologically valid measure of real-world perceptual categorization. Ratings showed substantial agreement across participants (speech; Fleiss κ = 0.69; music; Fleiss κ = 0.84). To relate these judgments to acoustic properties, we decomposed the audio signal using the STM framework (*46*), extracting STM patterns at each time point with a 1.5-second sliding window (see Figure 1B and Methods). This yielded time-resolved 2D matrices capturing spectral (0–9 cycles/kHz, sampled over 109 points) and temporal (0–15 Hz, sampled over 20 points) modulation power.

**Figure 1:**
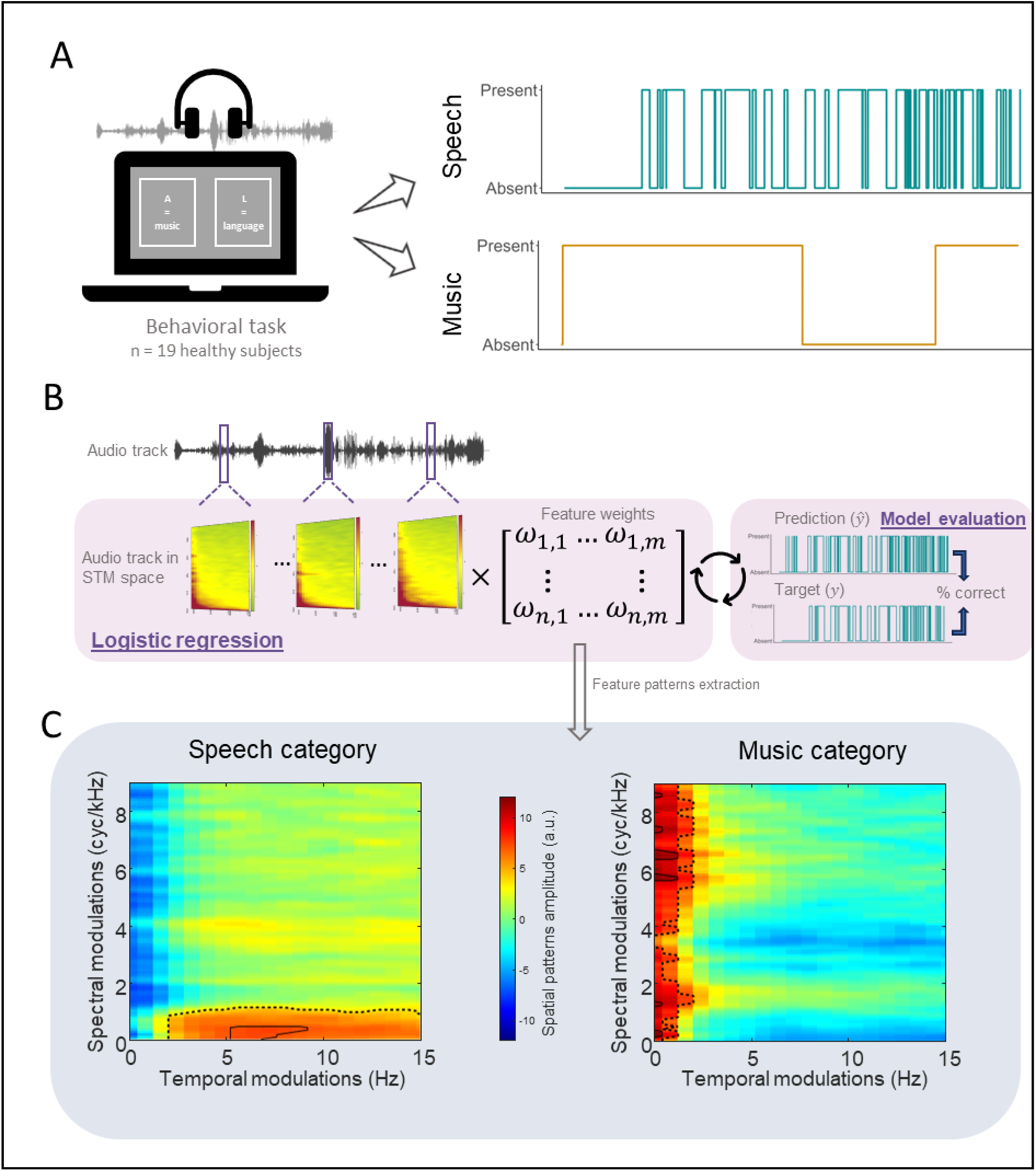
STM features predict perceptual categorization: spectral modulations for music, temporal modulations for speech. (A): Online behavioral task. Participants (n = 19) judged whether music and/or speech were present during the time-course of the audio track. The right panel illustrates judgement vectors obtained for speech and music categorization from one subject. Two judgement vectors were obtained for each participant. (B): Analysis pipeline. The audio track was decomposed into the STM space using a 1.5-second sliding window, generating a STM matrix for each time sample. For each category (speech/music), binary logistic regression models were trained on the audio track in the STM space to classify whether the corresponding category was present or absent along the time-course of the audio track. We chose to show an example model evaluation of the speech vector but note that independent logistic regressions for each category were performed to ensure that the feature contributions are fully interpretable and not influenced by joint optimization. The feature weights of the model were then extracted and transformed into interpretable feature patterns using the Haufe transformation (*47*). (C): Results. Averaged feature patterns across participants in the STM space for each categorical judgement. Threshold-based contours highlight regions of high feature patterns amplitude: the 90th percentile of the distribution is outlined with a dotted line, and the 99th percentile is outlined with a solid line.

Our first goal was to investigate for each of the behavioral participant if STM features are sufficient to explain categorization of speech and music, and if so which specific STM patterns contribute most to each category. We applied stratified 5-fold cross-validated logistic regression models to predict behavioral categorization based on the STM decomposition of the audio track. Each model used the STM data of all audio track samples as features to predict the binary behavioral judgment (presence/absence) outcome, for each category (speech or music). All 19 models showed a well-above-chance accuracy: 97-99 % for speech and 99-100% for music. Subsequently, we applied the Haufe transformation (*47*) to derive interpretable feature patterns from the feature weights, which were averaged for each judgement category across behavioral subjects (Figure 1C). For the speech category, the STM features most important for the binary classification mainly contained temporal modulations. Specifically, the 90th percentile of highest values spanned from 0.8 to 15 Hz in temporal modulations, but only from 0 to 1 cyc/kHz in spectral modulations (dotted line in Fig 1c, left). Peak activity (99th percentile; solid line in Fig 1c, left) was concentrated within 5.6 to 8.8 Hz in temporal modulations but 0 to 0.4 cyc/kHz in spectral modulations. For the music category, spectral modulations were instead the primary distinguishing features. High activity in the 90th percentile was observed across most of the spectral modulation range (except for a narrow gap between 3.3 and 3.7 cyc/kHz), and within a range of 0 to 1.6 Hz in temporal modulations (dotted line in Fig 1c, right). Peaks (99th percentile) were localized in a wide range of spectral modulation values, around 0.3, 1.2, 5.7, 6.5, 7.4, and 8.6 cyc/kHz, and temporal modulation range of 0 to 0.8 Hz (solid lines in Fig 1c, right). These findings align with the expected dynamics of high temporal/low spectral modulation for speech signals and low temporal/high spectral modulation for musical signals (*7*, *17*, *29*, *45*, *46*). Importantly, these results validate the use of time-resolved tracking in STM space to differentiate auditory categories. These findings were replicated in a complementary analysis using a fully balanced stimulus consisting of single-speaker speech and unrelated organ music that was entirely different from the Harry Potter soundtrack, and which provided us with ground-truth speech and music labels (see Complementary analysis in Supplementary materials and Figure S3A-B). For this data set models reached similarly high accuracy (∼99%) and revealed highly comparable feature patterns, with temporal modulations dominating speech classification and spectral modulations driving music classification. Building on this outcome, we next applied the STM tracking approach to assess which sEEG contacts in our patient sample could effectively track these STM features in the audio tracks that they listened to.

### Bilateral non-primary auditory areas track the spectrotemporal modulations of a naturalistic sound

Intracranial EEG recordings were collected from eleven patients with focal drug-resistant epilepsy as they passively listened to the audio track. After bipolar re-referencing, this resulted in a total of 846 sEEG contacts, with 51.4% located in the left hemisphere and 48.6% in the right hemisphere (Figure 2A, see Figure S1 for detail). For each sEEG contact, the magnitude of delta (0.5-3 Hz), theta (4-8 Hz), alpha (9-13 Hz), beta (14-25 Hz), low gamma (26-50 Hz), and high gamma (50-150 Hz) oscillations were extracted from the raw signal during the audio track listening, using Hilbert transform (z-score-normalized over time).

**Figure 2:**
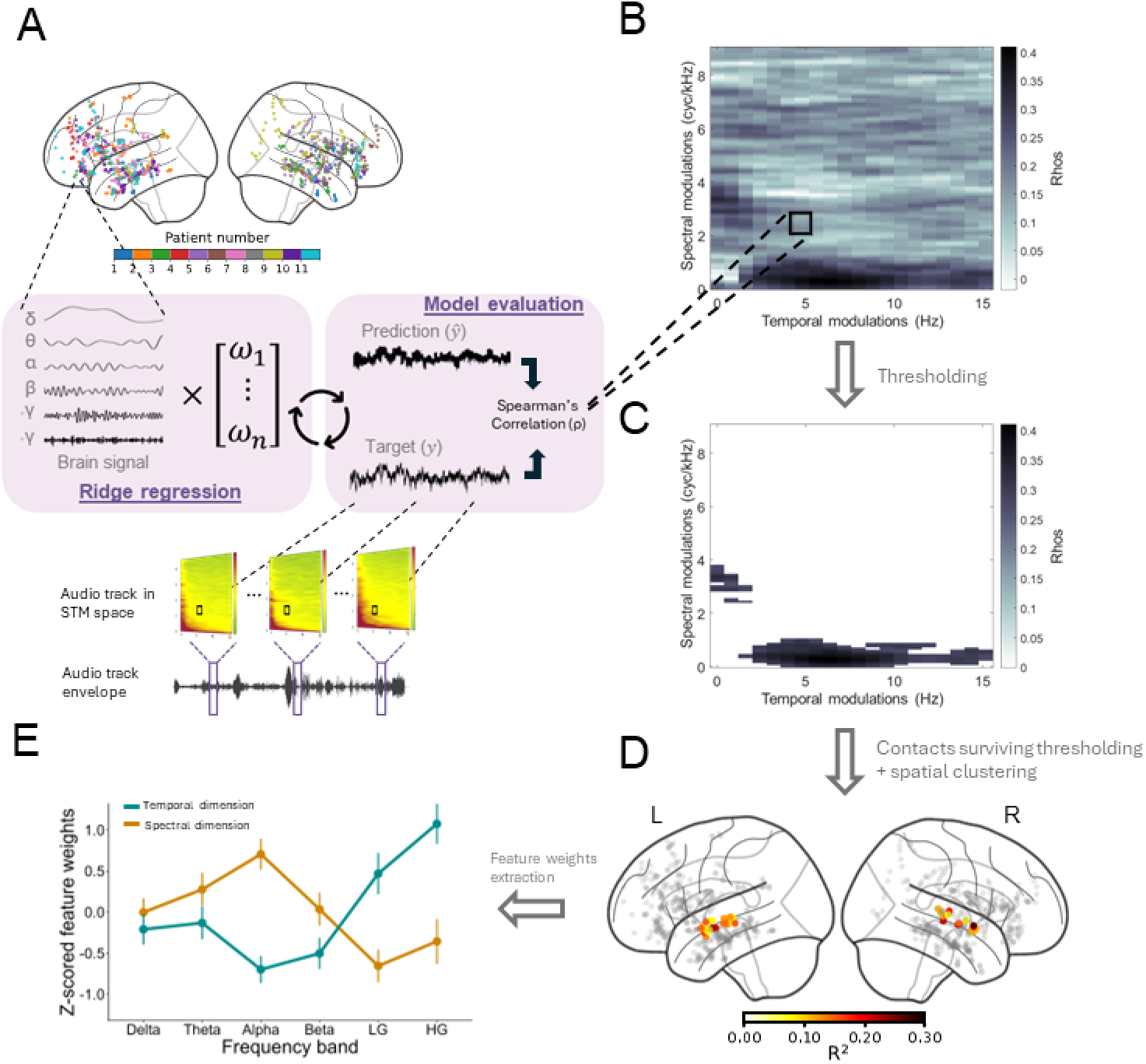
Bilateral non-primary auditory areas track the spectrotemporal modulations of a naturalistic sound. (A): Regression model schematics. sEEG contacts from eleven patients (top) with colors corresponding to each patient’s implantation scheme. Signal was decomposed into six frequency bands and used as features in a ridge regression model (middle). The audio track that the patients listened to (bottom part of panel a) was decomposed into the STM space as in Figure 1B. Each spectral/temporal combination time course from the matrix was then used as the target variable (*y*) in the regression model. Model predictions (ŷ) were evaluated (model evaluation panel) against the target (*y*) using Spearman’s correlations. (B): Example accuracy maps. An accuracy map was generated for each contact by running the model (depicted in (a)) at each spectral and temporal combination/coordinate in the STM space, yielding a map of correlation coefficient (ρ). (C): Thresholding. Each accuracy map was thresholded by performing, for each STM combination (n = 2180), a one-tailed one-sample t-tests against zero on Fisher z-transformed correlation coefficients. Correcting for the number of STM combinations, sEEG contacts and cross-validation folds yielded a Bonferroni threshold of approximately 2.7 × 10⁻⁹. We instead applied a more conservative fixed criterion of α = 10⁻³⁰, to retain only contacts showing robust STM tracking (see Methods). The heatmap represents the thresholded accuracy map from (B). (D): STM-sensitive contacts. Peak R^2^ values from contacts that survived statistical thresholding and spatial clustering (see Methods). Four clusters were identified in bilateral non-primary auditory areas (see Table S1 for coordinates and Figure S1 for detailed visualization and patients’ contribution). (E): Relative contribution of different frequency bands. Ridge model feature weights were extracted from the most informative spectrotemporal combinations (top 90th percentile identified via logistic regression masks, see Figure 1C) for speech (temporal dimension) and music (spectral dimension), z-scored across frequency bands, and averaged across selected combinations and time lags. Values are presented here averaged across STM-sensitive electrodes, with error bars representing standard error.

To uncover brain regions responsive to STMs, we applied 10-fold cross-validated ridge regression models that linked oscillatory magnitude across these six frequency bands, recorded from the 846 sEEG contacts, to the STM features of the audio track. For each sEEG contact, the model predicted the time course of STM amplitude for each of the 2,180 spectrotemporal combinations— corresponding to 109 spectral (0–9 cycles/kHz) × 20 temporal (0–15 Hz) modulation points extracted from the 2D STM matrices —producing an accuracy map of fisher-transformed correlation coefficients within the STM space. sEEG contacts that significantly predicted any portion of the STM space (t-tests vs 0, Bonferroni-corrected for number of STM points, sEEG contacts and cross-validation folds) were then spatially clustered in the MNI space, where clusters with overlapping contacts from at least three patients were considered significant (see (*48*) and Methods). Using this method, we identified four spatial clusters comprising a total of 39 sEEG contacts, bilaterally distributed in non-primary auditory cortical areas (see Table S1 for MNI coordinates and atlas labels). Two clusters were located in each hemisphere. In both hemispheres, one cluster included sEEG contacts primarily positioned in the anterior portion of the superior temporal region, with 11 sEEG contacts in the left hemisphere and 13 in the right. These anterior clusters included sEEG contacts located in the medial belt, in area A5, and the anterior distal superior temporal sulcus (STS), with the right cluster additionally containing an sEEG contact in area A4 according to the HCP atlas (*49*). The second cluster in the left hemisphere was located more posteriorly, encompassing sEEG contacts in the posterior distal STS and area A5. In the right hemisphere, the remaining cluster included sEEG contacts in the medial and lateral belt, and in area A5, corresponding to mid-to-posterior portions of the non-primary auditory cortex. Each cluster included sEEG contacts spanning at least three patients, ensuring the generalizability of these findings (see Table S1 for the MNI coordinates and atlas labels of the sEEG contacts). These results were replicated in a complementary analysis using the balanced speech and music stimulus described above, and an independent patient cohort (see Complementary analysis in Supplementary materials and Figure S3C-D), where a cluster of STM-sensitive electrodes was similarly identified in the right non-primary auditory cortex, despite a markedly different implantation scheme and limited left-hemisphere coverage. Overall, we identified sEEG contacts located in non-primary auditory areas that demonstrate the ability to track STM features of complex, naturalistic sounds. We then used the signal from these identified sEEG contacts for all subsequent analyses.

### Oscillatory frequency bands differentially support STM tracking

To quantify the relative contribution of each frequency band to STM tracking, we extracted z-scored feature weights from ridge models within STM-sensitive contacts, focusing on the most relevant spectrotemporal combinations (top 90th percentile; see Figure 1C) identified via binary masks from logistic regression patterns for speech (temporal dimension) and music (spectral dimension). This resulted in distinct measures of frequency-band contribution for each electrode and dimension (Figure 2E). A 6x2 Aligned Rank Transform test with frequency bands (delta; theta; alpha; beta; low-gamma; high-gamma) and dimensions (temporal; spectral) as within-subject factors revealed a significant main effect of frequency bands (*F*(5, 418) = 7.24, *p* < .001) and a significant frequency band-by-dimension interaction (*F*(5, 418) = 53.89, *p* < .001). Post-hoc analyses revealed a robust double dissociation: low- and high-gamma frequency bands made the strongest contribution to the prediction of temporal modulations (FDR-corrected *p* < .001 vs. all others) but made the weakest contribution to the spectral modulations (*p* < .02 vs. all others). Conversely, the alpha band, which dominated the spectral dimension (FDR-corrected p < .005 vs. all others), contributed the least to the temporal dimension (FDR-corrected *p* < .001 vs. all bands except beta, *p* = .17). Full post-hoc statistics are provided in Table S2.

### Behavioral categorization judgments can be decoded from brain-reconstructed STMs in bilateral non-primary auditory areas

We next address the key question of whether brain-reconstructed STMs (i.e., STMs predicted from neural signal) are sufficient to decode behavioral categorization judgments of speech and music. Our objective was to define how brain-reconstructed STMs can effectively link neural activity to behavioral judgment obtained in healthy individuals through acoustic features. At each time point, reconstructed STMs from STM-sensitive sEEG contacts (see Methods) were used as features to classify participants’ binary judgments of whether the track contained speech or music. Reconstructed STMs from 36 of the 39 STM-sensitive sEEG contacts significantly predicted speech presence (accuracy range: 54-60%; Wilcoxon signed-rank tests against zero across participants, FDR-corrected p < .046), while all 39 contacts significantly predicted music presence (54-71% accuracy values; FDR-corrected p < .001; see Figure S2). Consistent results were obtained in the complementary analysis using the balanced stimulus with ground-truth speech and music labels (see Complementary analysis in Supplementary materials and Figure S3F), where decoding was performed at the electrode level and reached similarly above-chance accuracy (53-86%) for both categories. These findings highlight that the category (speech or music) in audio tracks can be accurately predicted from brain activity, when solely represented in the STM space.

To identify the specific regions of the STM space contributing to category decoding across sEEG contacts, we performed, within each contact, a statistical test against zero on feature patterns at each point in the STM space across behavioral subjects (see Methods). We then counted, for each STM point, the number of contacts showing a significant feature pattern (Figure 3E). For the speech category, the STM regions with the highest number of sEEG contacts showing significant feature patterns correspond mainly to temporal modulations. Specifically, the top 10% of sEEG contact counts (90th percentile) ranged from 0.6 to 15 Hz in temporal modulations and from 0 to 1 cyc/kHz in spectral modulations (dotted line in Fig 3e, left). The peak sEEG contact count (99th percentile; solid line in Fig 3e, left) was concentrated in the range of 4.8 to 8 Hz for temporal modulations and 0 to 0.25 cyc/kHz for spectral modulations. For the music category, the STM regions with the highest number of sEEG contacts showing significant feature patterns corresponded mainly to spectral modulations. A high sEEG contact count in the 90th percentile was observed across most of the spectral modulation range, except for two narrow gaps between 3.4 and 3.7 cyc/kHz (consistent with the analysis using the actual audio track) and between 4.5 and 4.8 cyc/kHz (dotted lines in Fig 3e, right). The peak sEEG contact count (99th percentile; solid lines in Fig 3e, right) was localized within spectral modulation values around 0.2, 6, 6.6, 7.7, and 8.7 cyc/kHz and temporal modulation of 0 to 0.8 Hz. These results demonstrate that brain-reconstructed STMs can accurately predict behavioral categorization even though behavioral data were derived from different samples of participants (out of sample prediction). To assess the behavioral relevance of these brain-derived features, we will next test whether the STM patterns driving categorization are similar when derived from brain data (Figure 3E) and when derived directly from the acoustics (Figure 1C).

**Figure 3:**
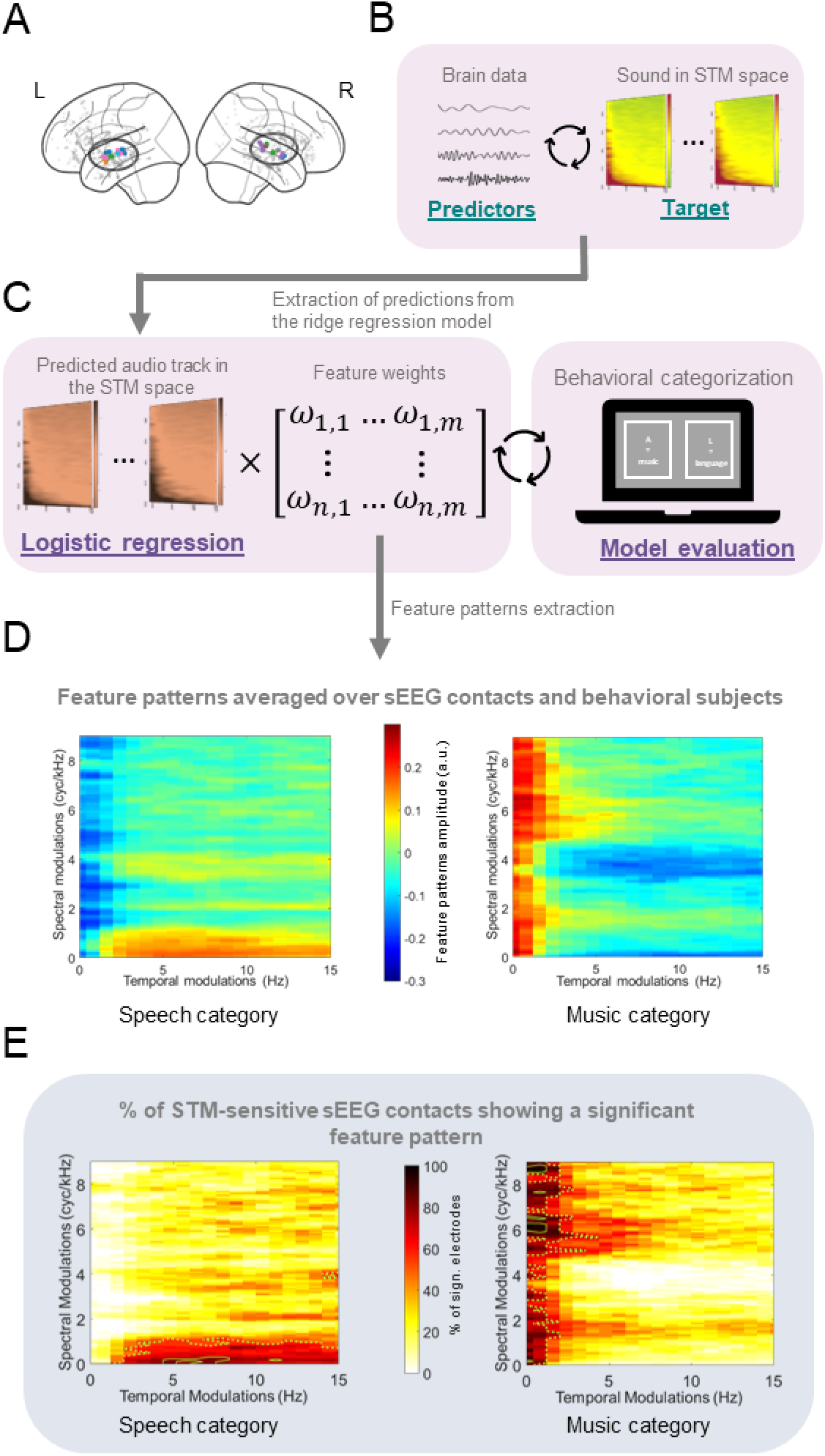
Behavioral categorization judgments can be decoded from brain-reconstructed STMs in bilateral non-primary auditory areas. (A) STM-sensitive sEEG contacts (same as Figure 2D). (B) Schematics of the regression model. The model used brain signals as predictors and the time courses of each spectral/temporal combination of the audio track as targets, as depicted in Figure 2A. The predicted time courses for each spectral/temporal combination were extracted, providing the audio track in the STM space predicted from brain signals. (C) Schematics of the logistic regression model. Based on the predicted audio track in the STM space obtained in (B), binary logistic regression models were trained to classify whether music or speech was present or absent (averaged accuracy values across participants for each electrode and category can be found in Figure S2). The feature weights of the model were then extracted and transformed into feature patterns using the Haufe transformation (*47*). Feature patterns were obtained for each STM-sensitive sEEG contact, behavioral participant and category. (D) Feature patterns in the STM space averaged over all sEEG contacts and all behavioral subjects. (E) Heatmaps showing, for each category, the percentage of STM-sensitive sEEG contacts that exhibited a significant feature pattern across behavioral subjects at each spectral and temporal combination/coordinate in the STM space. Threshold-based contours highlight regions of high sEEG contact count with significant feature patterns: the 90th percentile of the distribution is outlined with a dotted line, and the 99th percentile is outlined with a solid line.

### Brain-reconstructed and acoustic STM representations are highly similar

To assess whether the brain-reconstructed STM representations truly reflect the acoustic features relevant for perceptual categorization, we evaluated the similarity between the feature patterns (Figure 4A) of: (1) the STM of the actual audio track (Fig 1c), and (2) the brain- reconstructed STM (Fig 3d) averaged across hemispheres. For each category and each hemisphere, nineteen comparisons were performed between STM patterns derived from brain-reconstructed STMs and from acoustic STMs, both based on the prediction of each behavioral judgment vector (n = 19 behavioral participants). STM patterns were highly similar, showing significant correlation coefficients (Spearman correlation; p < .001, Bonferroni-corrected adjusted for number of subjects, categories, and hemispheres) for all comparisons, confirming the strong alignment between the STM features of the audio and those derived from brain activity (Figure 4B). We confirmed this result at the group-level, by comparing the distribution of correlation coefficients across behavioral participants to shuffled data, (see Methods; Figure 4B) revealing highly significant differences for each category (both *p* = 2.9x10^-22^). The complementary analysis using the balanced stimulus, also revealed that feature patterns derived from brain-reconstructed STMs were highly correlated with acoustic STM patterns across electrodes and at the group-level against a shuffled distribution (see Complementary analysis in Supplementary materials and Figure S3E). These key results demonstrate that the STM patterns in behaviorally relevant acoustic features are represented in a continuous and sufficiently similar manner in the brain in non-primary auditory areas. Next, we examined whether the representation of these STM patterns exhibits hemispheric asymmetries that have been previously associated with differential STM patterns processing.

**Figure 4:**
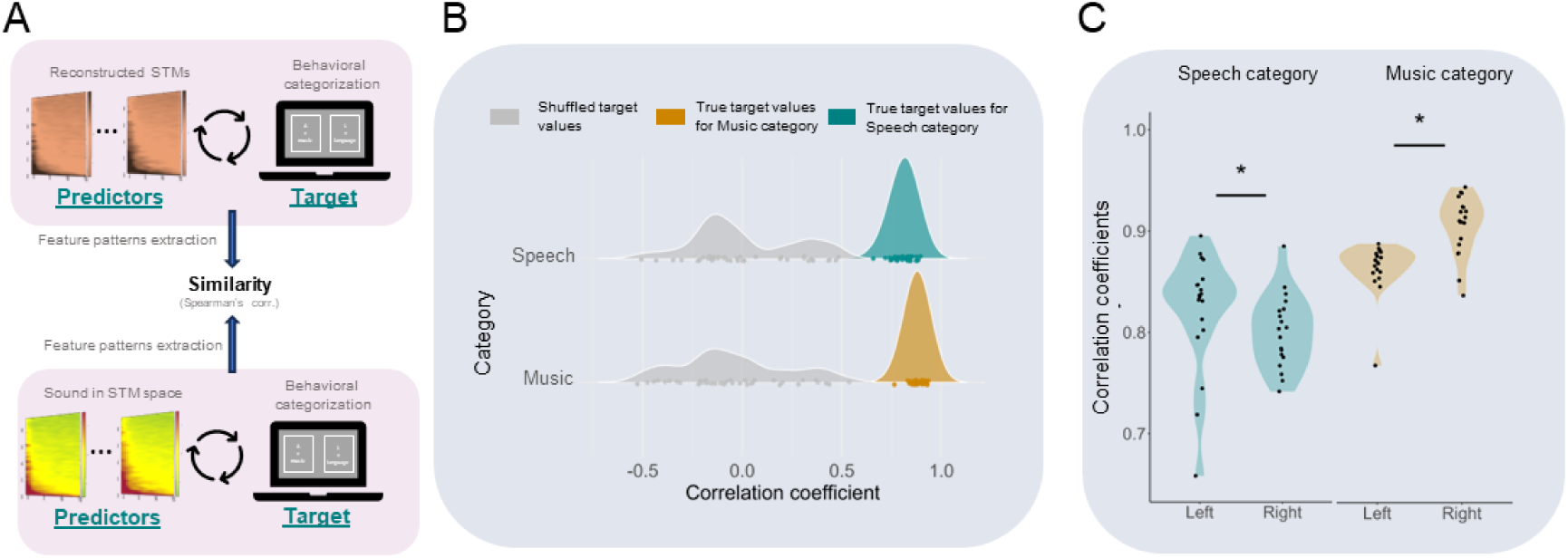
Similarity between brain-reconstructed and acoustic STM representations reveals hemispheric lateralization. (A) Schematic of the analysis pipeline. Feature patterns were extracted from the logistic regression models described in Figure 3C, where brain-reconstructed STMs served as predictors and behavioral categorization responses as target variables (top). This was done separately for each STM-sensitive sEEG contact, category, and behavioral participant. The resulting feature patterns were then averaged across sEEG contacts within each hemisphere. In parallel, feature patterns were obtained from the logistic regression models described in Figure 1B, where the actual acoustic STMs were used as predictors of behavioral categorization (bottom), for each category and behavioral participant. To evaluate similarity between brain-reconstructed and acoustic feature representations, Spearman correlations were computed between the two sets of feature patterns. For each category and each hemisphere, this resulted in 19 comparisons (one per behavioral participant) between the hemisphere-averaged brain-reconstructed STM patterns and the corresponding acoustic STM patterns. (B) Feature pattern similarity distribution. Density curves represent the distribution of correlation coefficients between feature patterns derived from the two models depicted in (A). The top (aqua) distribution corresponds to the speech category, and the bottom (orange) distribution corresponds to the music category. For comparison, a separate distribution (gray) shows the correlation coefficients obtained when feature patterns from the first model (top panel of (A)) were compared with those from the second model after shuffling categorization labels. All Spearman’s correlation coefficients calculated from the two original models were significant at *p* < .001 after Bonferroni correction adjusted for number of subjects, categories, and hemispheres. (C) Similarity lateralization. Correlation coefficients are displayed for each behavioral participant in the speech category (left) and the music category (right), with data from sEEG contacts implanted in the left and right hemisphere (x-axis). \**p* < .01, with pairwise Wilcoxon signed-rank tests, FDR-corrected.

### Similarity between brain-reconstructed and acoustic STM representations reveals hemispheric lateralization

Finally, we tested whether the similarity between the two representations reflected the previously described hemispheric specialization of the auditory cortices (Figure 4C), with the left hemisphere being more sensitive to temporal modulations and the right to spectral modulations. To do so, we carried out a 2x2 aligned rank transform test on the correlation coefficients used in the previous similarity analysis, with category (speech; music) and hemisphere (left; right) as factors. The main effect of category was significant (*F*(1, 55.65) = 72.20, *p* < .001), with higher overall correlation coefficients for the music category compared to the speech category. No main effect of hemisphere was observed (*F*(1, 55.65) = 0.68, *p* = 0.412), indicating that both hemispheres similarly encode auditory categorical information. Finally, the category-by-hemisphere interaction was significant (*F*(1, 55.65) = 23.64, *p* < .001). Post-hoc analysis showed that for the speech category, correlation coefficients were significantly higher in the left hemisphere (FDR-corrected *p* = .027) compared to the right, whereas for the music category, correlation coefficients were significantly higher in the right hemisphere (FDR-corrected *p* < .001) compared to the left. Overall, we found that the high similarity between feature patterns derived from acoustical-STM and brain-reconstructed STM showed hemispheric asymmetry with greater similarity in the left hemisphere for speech categorization and in the right hemisphere for music categorization.

## DISCUSSION

In the present study, we investigated how the human brain encodes and categorizes naturalistic speech and music, integrating independent continuous behavioral categorization judgments from healthy participants with intracranial recordings from the auditory cortex of epileptic patients. We specifically tested whether the cortical representation of acoustical features carries sufficient information to account for the perceptual categorization of speech and music. Our results converge on the conclusion that the spectrotemporal modulation (STM) features of naturalistic sounds, as they are represented in non-primary auditory cortex, are sufficient to support the categorization of speech and music. Specifically, and in line with our stated predictions, we found that (i) distinct STM patterns are associated with the categorization of naturalistic speech and music categories in healthy individuals (Figure 1C), validating our analysis approach to naturalistic auditory input; (ii) STM features are robustly tracked in non-primary auditory areas (Figure 2D); (iii) STM features are encoded with a structured oscillatory code, such that alpha-band activity preferentially supports the encoding of spectral modulations whereas low- and high-gamma activity preferentially supports the encoding of temporal modulations, a double dissociation that identifies separable oscillatory mechanisms for the two dimensions of the STM representation (Figure 2E); (iv) STMs reconstructed from brain signals in these regions are sufficient to predict behavioral categorization judgments from a separate group of participants (Figure 3E), showing that brain-derived representations align closely with perceptual decisions; (v) STM patterns from the brain-reconstructed STMs are highly similar to those obtained from the actual audio track (Figure 4B), and (vi), this effect is lateralized (Figure 4C) with stronger acoustic/brain similarity in the right hemisphere for music and in the left hemisphere for speech, consistent with well-known hemispheric asymmetries in auditory processing. Importantly, the behavioral, tracking, decoding and similarity findings were fully replicated in a complementary dataset using a distinct set of patients and stimuli (Complementary analysis and Figure S3E), strengthening the generalizability of our results. The hemispheric asymmetry could not be assessed in that dataset, in which electrode coverage was restricted to the right hemisphere, and the oscillatory dissociation was not recovered, the band-resolved estimates being less stable from a single cluster than in the four bilateral clusters of the main analysis. We next discuss each of these points in turn.

### Distinct STM patterns are associated with the perception of naturalistic speech and music categories

Using a time-resolved tracking approach, we found that human judgments of perceiving speech and/or music at any given moment within a naturalistic soundtrack can be well differentiated based on their STM content, with speech associated with high temporal (peak 5.6–8.8 Hz) and low spectral modulations (0–1 cyc/kHz), and music with high spectral (peaks between 5.7 and 8.6 cyc/kHz) and low temporal modulations (0–1 Hz , Figure 1C). This finding is consistent with prior literature demonstrating that speech and music rely on distinct STM features, which not only support their perceptual differentiation but also provide an optimal representational framework for capturing their complex structures. Specifically, temporal modulations in the range of 4–7 Hz are known to be critical for speech intelligibility, as this range aligns with the syllabic rate of spoken language (*17*, *46*, *50–52*). In the spectral domain, speech recognition has been shown to rely primarily on low spectral modulations (*17*), with critical modulations for speech comprehension typically falling below 4 cyc/kHz (*46*). In contrast, music perception operates on a different timescale, with key temporal modulations typically falling below 2 Hz (*53*), reflecting the slower amplitude modulations of musical phrases across genres (*50*) and cultures (*17*). Regarding spectral modulations, previous studies have shown that for songs that contain only vocal elements, important spectral modulations range from approximately 3 to 8 cyc/kHz (*7*, *17*). Our present results and previous literature thus confirm that STM representations can effectively capture the statistical properties of speech and music that are most relevant for perceptual categorization.

These results were obtained under conditions that differ in an important respect from those in which the STM framework has previously been evaluated. In the present soundtrack, speech and music alternate and frequently overlap, and in the complementary stimulus set used for replication, speech-only, music-only and overlapping segments each account for a third of the total duration, so that the models had to determine the presence of each category while the other was often present as well, rather than to discriminate between two classes of sounds presented in isolation. The fact that STM representations support categorization under these conditions indicates that the acoustic distinction between speech and music remains available within a continuously varying mixture of the kind encountered in everyday listening, and not only when the two are compared as separate and acoustically homogeneous stimuli. These results next raise the question of whether such behaviorally relevant acoustic representations are also reflected in the tuning properties of the human auditory cortex.

### The auditory cortex robustly tracks STMs

Our second finding demonstrated that the auditory cortex is tuned to track the STM regularities present in natural speech and music. We identified sEEG contacts in non-primary auditory cortex that significantly tracked STM patterns, using a robust ridge regression approach combined with the temporal response function method (Figure 2A). We demonstrated that STM patterns are dynamically tracked in non-primary auditory regions (Figure 2D). Specifically, STM tracking was observed in comparable locations across both hemispheres in the medial and lateral belt regions, area A5 and the distal superior temporal sulcus (STS). No such tracking was observed in sEEG contacts outside these auditory regions. Importantly, none of our participants had sEEG contact coverage over the primary auditory cortex, a common limitation of intracranial studies, so we cannot exclude the possibility that STM tracking may also occur in primary or adjacent auditory areas.

Our findings align with a broad body of evidence from both animal and human studies showing that STM sensitivity is a fundamental property of the auditory system. In animal models, single-unit recordings across species such as zebra finches, ferrets, cats, and monkeys have consistently revealed tuning to STM throughout the auditory pathway. Tuning to STM has been reported from the inferior colliculus and medial geniculate body to primary and secondary auditory cortex (primarily using methods that estimate spectrotemporal receptive fields (STRFs) of individual neurons) in response to controlled synthetic ripple stimuli as well as more ecologically valid sounds like birdsong and vocalizations (*8*, *10*, *15*, *16*, *37*, *54–57*). Similarly, human neuroimaging research has confirmed STM sensitivity across primary, secondary, and even higher-order auditory regions. This has been demonstrated through multiple approaches: spectrotemporal modulation transfer functions measured with synthetic ripple stimuli (*12*), STRFs estimated with naturalistic sounds (*39*, *58*), classic activation-based fMRI mapping using dynamic ripple stimuli (*59*), and encoding/decoding models that explicitly linked neural responses to the STM content of speech, music, vocalizations, and environmental sounds (*11*).

Further evidence for STM tuning has been shown recently in three human studies conducted within the STM framework. These studies demonstrated that degrading the STM of speech and tonal sounds leads to reduced cortical discriminability, with effects localized in the same regions as our STM-sensitive contacts. Albouy et al.(*7*) showed in an fMRI study that STM degradation of sentences and melodies impaired machine-learning based category classification in left A4 and medial belt for temporal modulations, and right medial belt for spectral ones. Robert et al. (*31*) used synthetic sounds mimicking simple actions (containing mainly temporal modulations) and object properties (containing mainly spectral modulations) and found that STM degradation disrupted fMRI-based classification in left A4/A5 and IFG for actions and right parabelt, A4/A5 and pSTS for objects. Flinker et al.(*45*), using electrocorticography and degraded speech, reported cortical tracking of STM features in left A5 (temporal) and right medial belt (spectral) for speech- and vocal timbre-related features respectively. Across these studies, despite differences in stimuli and methods, sensitivity to STM structure consistently emerged in areas overlapping with the STM-sensitive contacts identified in the present study.

A different interpretation has been proposed based on fMRI voxel decomposition methods, according to which speech and music categories are associated with neural components that are distinct from the components sensitive to acoustic features, and whose selectivity was not accounted for by the STM features against which it was evaluated (*21*, *22*). Yet, the topographies they report largely overlap with the STM-sensitive regions identified in the present and previous studies. In that fMRI work (*21*), independent components tuned to STMs were located near the borders of primary auditory cortex, both anteriorly and posteriorly, while speech- and music-selective category components emerged in lateral STG, planum polare, and planum temporale. Similarly, in an electrocorticography (ECoG) study (*22*), the authors reported STM sensitivity in both primary auditory cortex and more anterior non-primary auditory areas and revealed speech- and music-selective components in middle and anterior STG, in proximity to the clusters found in the present study (Figure 2D). Thus, despite interpretative differences, the findings across studies converge regarding both the existence and the location of higher-order representations for speech and music, a point we shall return to below. However, it should be noted that the studies using voxel decomposition do not show hemispheric differences, unlike the findings reported here.

### Oscillatory frequency bands differentially support STM tracking

Building on the identification of STM-sensitive electrodes, we next investigated whether STM tracking for speech and music relies on a structured oscillatory code. Given the established principled relationship between the time scales present in complex sounds and the time constants of cortical oscillations (*42*), we hypothesized that this framework would extend to spectrotemporal modulations, supporting distinct oscillatory contributions to speech- and music-relevant STM dimensions. We observed a clear double dissociation: low- and high-gamma bands contributed most to the prediction of temporal modulations, which are critical for speech, whereas alpha contributed most to spectral modulations, which are more informative for music (Figure 2E). These results are consistent with intracranial evidence that primary and secondary auditory regions display bimodal response profiles with peaks in the theta and beta/gamma ranges, in line with a two-timescale processing mode, and that responses in association auditory cortex become more heterogeneous for complex stimuli, with alpha and gamma activity co-occurring bilaterally (*43*). Because the STM-sensitive contacts identified here were located in non-primary auditory regions, the joint contribution of alpha and gamma activity that we report is in agreement with this observation and extends it by showing that the multiplexing of neural timescales is organized along the two dimensions of the STM representation, with distinct frequency bands preferentially carrying the temporal and the spectral modulation content of the same acoustic signal.

The double dissociation reported here does not imply an alignment between the frequency of the neural rhythm and the frequency of the acoustic modulation, since the temporal modulations analyzed span 0 to 15 Hz and the oscillatory bands were included in the model through the amplitude of band-limited activity rather than through their phase. It does, however, place the two dimensions of the STM representation on different neural timescales, which can be related to the proposal that the duration of an oscillatory cycle sets the length of the window over which the acoustic signal is analyzed (*42*, *43*). Under this proposal, the shorter cycles of the gamma range would provide the resolution required to follow rapid fluctuations along the temporal axis, whereas the longer cycles of the alpha range would provide the extended integration required to resolve fine detail along the spectral axis, these two forms of resolution being traded against one another for any given window length. This interpretation is consistent with the framework within which auditory hemispheric asymmetries have been described (*7*, *33*, *43*), but it remains to be tested directly, since the present analyses quantify the contribution of the amplitude of each frequency band rather than the windows over which the signal is integrated.

### Brain-reconstructed STMs are sufficient to decode behavioral categorization judgments

Having established that STMs provide an efficient shared acoustic representation for distinguishing speech and music, and that the auditory cortex is finely tuned to these features via frequency-specific oscillatory contributions to STM tracking, we reconstructed the STM features of the audio track from intracranial signals recorded at STM-sensitive sEEG contacts (Figure 3A-C). We found that the predicted STMs were highly effective in decoding continuous categorization judgments obtained out-of-sample. This result suggests that behavioral categorical decisions can be predicted from brain-reconstructed acoustic representations. In addition, a closer look at the feature patterns of the brain-reconstructed STMs that were most predictive of behavioral categorization (Figure 3D) revealed a match with those identified using the STM patterns derived directly from the real audio track (Figure 3E). This observation suggests a functional convergence between acoustically and neurally derived features. The next important step was therefore to formally test the similarity between these two sets of STM patterns to determine whether the same representational structure underlies both acoustically and neurally derived data.

### High similarity between brain-reconstructed and acoustically derived STMs

A similarity analysis (Figure 4A) revealed a strong correspondence between the acoustically and neurally derived STM patterns that are critical for successful behavioral categorization (Figure 4B). Notably, because our neural and behavioral data were collected from separate participant groups, the observed neural–perceptual mapping reflects a robust out-of-sample prediction, a rigorous validation approach in cognitive neuroscience (*60*), indicating that the underlying coding principles generalize across individuals rather than overfitting to subject-specific variability or noise. These findings therefore strongly suggest that brain-reconstructed STM representations alone are sufficient to accurately predict behavioral categorization judgments of speech and music, establishing a link between neural encoding of STM features in the auditory cortex and higher-order perceptual outcomes. The final analytical step focused on whether this acoustic-neural coupling reflects meaningful organizational principles of auditory processing, by testing for hemispheric specialization.

### Similarity between brain-reconstructed and acoustic STM representations reveals hemispheric lateralization

Hemispheric asymmetries (favoring the left auditory cortex for speech and the right for music) have been robustly documented over decades of research, including lesion studies, electrophysiological recordings, and neuroimaging investigations in both neurotypical individuals and those with neurodevelopmental disorders such as amusia (*61–66*). Our data revealed a hemispheric lateralization in the similarity between brain-reconstructed STM patterns (extracted from STM-sensitive sEEG contacts) and the STM patterns of the actual audio stimuli, when both were used to predict perceptual categorization outcomes (Figure 4C). Specifically, speech categorization showed higher similarity in sEEG contacts implanted in the left-hemisphere, while music categorization showed higher similarity in sEEG contacts implanted in the right-hemisphere. Our results thus show that the brain leverages the statistical structure of STM in a category- and hemisphere-specific manner to support auditory categorization.

In line with our results, hemispheric specialization for STM processing has been well-documented: the left auditory cortex operates at faster temporal integration windows than the right (*28*, *43*, *67*), making it better suited for tracking rapid temporal modulations, such as those prominent in speech. Conversely, the right auditory cortex exhibits sharper spectral tuning and heightened sensitivity to pitch-related information (*33*, *68–71*), aligning with its preferential processing of fine spectral cues relevant for music. More recently, this hypothesis has been directly tested using the STM framework across fMRI, MEG, and sEEG/ECoG, showing that degradation of speech, songs, or environmental sounds in the temporal or spectral modulation domains selectively impact brain activity in the left STG for temporal degradation and in the right STG for spectral degradation (*7*, *31*, *45*). The present findings and recent literature reinforce the view that auditory perception relies on a shared STM encoding framework, with category-related hemispheric biases emerging as a consequence of distinct sensitivity in the processing of temporal and spectral modulations rather than as a function of intrinsic domain specificity.

### Efficient neural coding in the auditory system

Our results can be understood within the broader efficient neural coding framework, which posits that perceptual systems have evolved to efficiently encode behaviorally relevant environmental stimuli for any given species (*1*). Computationally, efficient coding can be achieved if neural representations match the statistical regularities of the signals they represent (*72*). While this principle has been extensively documented in the visual domain (*73*, *74*), it has also been increasingly supported in the auditory modality (*75–78*). Among various ways to represent natural sounds, STMs are particularly well-suited for capturing the statistical structure of complex sounds (Figure 1C), including vocalizations, as they reflect joint dependencies between temporal and spectral modulations that are characteristic of natural stimuli, dependencies that are not captured by the raw spectrum or spectrogram alone (*5*, *13*, *14*).

At the neural level, STMs capture auditory response properties beyond traditional frequency tuning across species and regions, including the midbrain, thalamus, and primary/secondary cortices of birds and non-human mammals (*8*, *15*, *16*, *37*, *56*). In humans, specific STM tuning in the auditory cortex (including non-primary regions) has been demonstrated via neuroimaging with synthetic dynamic ripples (*12*, *59*) and natural speech (*11*, *39*, *58*), as well as through intracranial recordings using speech (*38*). In addition, speech signals can be reconstructed from patterns of activity in the STG using STM-based STRF models (*9*). Accordingly, under the assumption of efficient neural coding, and given the known tuning of central auditory neurons to STMs, we propose that category-specific neural representations in the auditory cortex, such as those distinguishing speech from music, can be largely explained by differential sensitivity to STM features. It should be made explicit that the present design does not isolate category-selective representations that would be independent of STM coding. Naturalistic speech and music preserve a substantial covariance between acoustic structure and category identity, and the neural representation used to predict behavior was restricted to STM features, so that the analyses reported here establish the sufficiency of STM coding for category-level differentiation rather than the absence of additional categorical representations.

While we argue that STM representations are sufficient to support category-level distinctions, we do not dispute the likely existence of subsequent transformations and additional complex processing that may support higher-order, abstract perceptual processes. Specifically, even if categorical representations depend on characteristic low-level features, the specific content of these representations would require much more fine-grained analysis that goes beyond the spectrotemporal distribution, in order to identify and encode particular items within categories. A critical consideration in this context is the hierarchical nature of auditory cortical organization (cf. (*19*)), as supported by extensive evidence in both human and non-human primates (*6*, *79*), paralleling the functional architecture observed in the visual system (*80*, *81*). From this perspective, category-selective responses observed in non-primary auditory cortex through non-linear voxel decomposition methods (e.g., (*21*, *22*)) may reflect higher-order transformations that build on, rather than replace, STM-based representations. Notably, the brain regions identified in those studies as selectively responsive to speech or music largely overlap with the STM-sensitive regions revealed in our present work and in previous reports, with the additional property that they are lateralized. This overlap suggests that the emergence of categorical selectivity in these areas may be a consequence of their differential sensitivity to the statistical structure of specific sound categories, as captured by STMs. In this view, the observed selectivity is not at odds with a domain-general encoding scheme but rather emerges naturally from it. Selectivity that emerges from differential sensitivity to STM structure does not need to account for every category-selective response, however. It has been shown, for instance, that music-selective responses in the superior temporal gyrus track the statistical expectation of upcoming notes, with a corresponding sensitivity to phoneme-level expectation at speech-selective sites (*40*). Properties defined over the sequence rather than over the momentary spectrotemporal content illustrate the type of higher-order transformation that would build on, rather than replace, the acoustic representation characterized here.

In conclusion, our findings that neural STM representations efficiently capture the statistical properties of naturalistic speech and music thus can be viewed in the context of functional adaptation of the human brain to these important classes of sounds. The present study contributes to a longstanding theoretical question by showing that efficient neural coding of STMs provides a biologically plausible framework that is sufficient to support category-level distinctions, both at the neural and behavioral levels. It further shows that the temporal and the spectral components of this representation are carried by distinct oscillatory mechanisms, and that the correspondence between brain-reconstructed and acoustically derived STM patterns is lateralized, being stronger in the left hemisphere for speech and in the right hemisphere for music. While our findings do not exclude category-specific mechanisms at later stages of processing in higher-order associative brain regions, they indicate that the acoustic domain already provides the information required for these distinctions.

Finally, it is important to recall that the auditory processing hierarchy does not start at the cortical level. Subcortical structures such as the inferior colliculus and the medial geniculate body exhibit tuning to spectrotemporal features in animal models (e.g., (*15*, *16*)). Investigating these subcortical contributions in humans could provide valuable insights into the early stages of hierarchical auditory encoding and help refine our understanding of how STM representations emerge and propagate along the auditory pathway.

## MATERIAL AND METHODS

### Patients

sEEG data from 11 patients suffering from drug-resistant focal epilepsy were collected (self-reported biological sex, 4 females, mean age = 29.6 years, SD = 3.2 years) following the Research Opportunities in Humans Consortium guidelines (*82*). All patients were native French speakers from Canada and they all completed the task as instructed. The study protocol was explained verbally by both the research and clinical teams, and participation was discussed with each patient prior to hospitalization. All patients provided verbal assent and signed written informed consent forms. This research complies with all relevant ethical regulations; the experimental procedures were approved by the Ethics Review Board of the CHU de Québec - Université Laval (2022-5890, Québec, QC, Canada).

### Healthy Participants

20 healthy adults participated in the behavioral experiment online. No statistical method was used to predetermine sample size. All participants were native French speakers from France or Canada (self-reported biological sex, 11 females, mean age = 36.4 years, SD = 14.7 years). Participants reported normal hearing. Data from one participant was excluded due to a problem with data collection on the online server. All participants provided written informed consent, and the experimental procedures were approved by the Ethics Review Board of the CIUSSS de la Capitale Nationale (2022–2476). The study has been conducted according to the principles expressed in the Declaration of Helsinki.

### Stimuli and procedure

#### Stimuli

The stimuli used in both the sEEG and behavioral experiments were identical and consisted of the original audio track from the first 7 minutes and 50 seconds of the French-dubbed version of *Harry Potter and the Philosopher’s Stone* (titled *Harry Potter à l’école des sorciers* in French). This audio track included the complete soundtrack from the movie, including speech, music, and environmental sounds as they appear in the cinematic experience.

#### Procedure: sEEG experiment

sEEG patients, who had electrodes implanted based on clinical requirements, were individually tested in a dedicated sEEG monitoring room at the *Hôpital de l’Enfant Jésus*, CHU de Québec (QC, Canada). sEEG data were recorded continuously throughout the session. Stimuli were presented using Presentation software (Neurobehavioral Systems, Albany, CA, USA). Patients were given the option to use their own earphones or those provided by the research team. After ensuring a comfortable sound level with a separate audio file, patients were instructed to listen attentively to the entire audio track. Behavioral categorization judgements were not acquired in patients as this procedure aimed at capturing brain activity under ecologically valid conditions, where listeners are not actively categorizing or labeling stimuli (e.g., “this is speech,” “this is music”), as such cognitive demands could introduce additional neural activity unrelated to passive perception.

#### Procedure: Behavioral experiment

Participants completed a judgment task to determine whether music or speech was present in an audio track. The experiment was designed using PsychoPy (*83*) and hosted on Pavlovia for online data collection (data available at https://doi.org/10.17605/OSF.IO/4726S). Upon accessing the experiment, participants provided demographic information, including age and biological sex. They were instructed to ensure they would be undisturbed for at least 15 minutes, wear personal headphones, and adjust the volume to a comfortable level using an audio excerpt. The task instructions explained that participants were to press the "Q" key whenever they heard music and the "L" key whenever they heard speech, holding the key as long as the respective category was present and releasing it when it was absent. They were explicitly instructed that both keypresses could happen at the same time, thus making the judgement task simultaneous for both categories. A visual feedback screen displayed two rectangles labeled "Q = Music" and "L = Language" (see Figure 1A) which turned red when the corresponding key was pressed and reverted to white when released. Prior to starting, participants were asked to try pressing both keys to familiarize themselves with the feedback. They then completed a 2-minute training session using an unrelated audio track with both music and conversation before proceeding to the main task, which involved the 7-minute-and- 50-second experimental audio stimuli described earlier.

### Extraction of spectrotemporal modulations

The acoustic signal was analyzed over its time course using the STM framework (see *7*, *17*, *46* for similar procedures). STM patterns (i.e., modulograms) were extracted to obtain 512 STM per second, a rate corresponding to the sEEG sampling frequency using the ModFilter algorithm (*46*) with a 1.5-second window starting from the current time sample (see Figure 1B). For each window, this method applies a 2D fast Fourier transform to the autocorrelation matrix of the sound stimulus in its spectrographic representation. This process generates, at every time sample, a 2D matrix capturing the power of the audio signal across spectral and temporal combinations. The STM pattern extraction was performed separately for each channel of the stereo audio track, and the resulting patterns were averaged together in the STM space to obtain a unified representation of the audio stimulus. Because speech and music STM patterns are typically symmetric between positive and negative temporal modulations (*46*) and to reduce data size for faster computation, the negative temporal modulation values were averaged with the corresponding positive values. This resulted in STM patterns at each time sample, with temporal modulations ranging from 0 to 15 Hz (20 points) and spectral modulations spanning 0 to 9 cycles/kHz (109 points), corresponding to a total of 2180 spectrotemporal combinations.

### Behavioral data

For each participant and each category (speech/music), a binary judgement vector was created, with 1 indicating that the participant perceived the category in the audio track, and 0 indicating its absence. Inter-rater reliability for the binary coding of speech/music presence across the time course of the audio track was assessed using Fleiss’ Kappa (κ, *84*). This statistic assesses the degree of agreement between more than two raters (here, N=19) assigning categorical ratings (0 = absent, 1 = present) to a fixed set of items (here time points). Analyses were conducted separately for the speech and music categories using the *irr* package in R (*85*).

### sEEG recording and preprocessing

#### sEEG contacts localisation

Patients were stereotactically implanted with 16 to 18 semi-rigid multi-lead electrodes (AdTech; diameter: 0.8 mm). Brain activity was recorded from sEEG contacts, which were 2.0 mm wide and spaced 3, 4, 5, or 6 mm apart, targeting clinically defined brain structures. A 3D T1-weighted anatomical MRI (MPRAGE, 3T Siemens Trio, Siemens AG, Erlangen, Germany) was acquired pre-implantation, consisting of 160 sagittal slices with 1 mm³ isotropic voxels covering the entire brain. Cortical surfaces were reconstructed from the T1-weighted MRI, and the precise locations of the sEEG contact were identified using a post-implantation CT scan co-registered to the pre-implantation MRI (ImaGIN toolbox, https://f-tract.eu/software/imagin/). The MNI coordinates of the sEEG contacts were determined using the SPM toolbox (http://www.fil.ion.ucl.ac.uk/spm/). Finally, atlas labels were extracted from the extended Human Connectome Project (HCPex) multimodal parcellation atlas (*86*), a modified and expanded version of the HCP-MMP v1.0 atlas (*49*).

#### Recording

The sEEG data were acquired with a hardware band-pass filter of 0.1–200 Hz with a sampling rate of 512 Hz (Nyquist frequency of 256 Hz) using a Natus video-sEEG monitoring system. This configuration provides a sufficient buffer to investigate the high-gamma band up to 200 Hz without significant attenuation or aliasing. Online referencing was performed using a contact located in white matter. During acquisition, signals were bandpass filtered between 0.1 and 200 Hz. sEEG contacts contaminated by pathological epileptic activity or environmental artifacts were rejected in collaboration with the clinical team.

#### Preprocessing

The Brainstorm software with default parameters (*87*) was used to preprocess and Hilbert transform sEEG data. A notch filter was applied to reduce powerline interference (60 Hz and its harmonics, 120 and 160 Hz). The sEEG signal was segmented from -10 seconds to 490 seconds relative to the audio track onset, (-10 to 0 seconds baseline period). Each sEEG signal was subsequently re-referenced to its immediate neighbor using bipolar derivations. This bipolar montage enhances local specificity by removing artifacts common to adjacent contacts, such as powerline noise (e.g., 60 Hz) and distant physiological artifacts, while also reducing the effects of distant sources propagating through volume conduction. The resulting spatial resolution of each sEEG bipolar contact is estimated at 3–4 mm (*88*). Brain activity was recorded from a total of 846 bipolar contacts (ranging from 47 to 88 across patients).

### Hilbert transform

To investigate the relationship between oscillatory rhythm fluctuations and STM in the audio track, we applied the Hilbert transform to extract the amplitude envelope (i.e., magnitude) of oscillatory rhythms across the entire neurophysiological range. This analysis was performed for each sEEG contact during the audio track listening period. Oscillatory rhythms were segmented into the following frequency bands: delta (0.5–3 Hz), theta (4–8 Hz), alpha (9–13 Hz), beta (14–25 Hz), low gamma (26–50 Hz), and high gamma (50–150 Hz).

#### Quantification and statistical analysis

The logistic regression analyses were conducted with the *MNE-Python* (*89*) and *Scikit-learn* (*90*) libraries.

The ridge regression analyses were conducted with the *spyeeg* library (*91*). Statistical analyses were conducted with MATLAB Version: 9.12.0 (R2022a).

Spatial clustering was conducted with custom-made MATLAB scripts (https://doi.org/10.17605/OSF.IO/4726S).

### STM features used in perceptual categorisation

For each behavioral subject and each category (speech/music), a logistic regression model was applied, where the STM of all samples of the audio track served as features, and the binary behavioral judgment of the category acted as the target variable. We chose to fit an independent logistic regression model for each category to ensure that the feature weights for speech and music were entirely independent and directly interpretable. We avoided joint or dual logistic regression to prevent any potential interactions between outputs that could bias the weights. Both the features and target variables were temporally downsampled to 50 Hz to optimize processing efficiency. The data were shuffled before splitting into folds to ensure representative and unbiased partitions, and the model was evaluated using stratified 5-fold cross-validation. This combined approach ensured robust generalization and mitigated potential biases from the data’s original ordering. To verify that all models were successfully trained, we computed the classifier’s performance as the mean accuracy across the 5 stratified cross-validation folds. Model accuracies for each category were significantly above chance across subjects (all adjusted p < .001), as determined by a Wilcoxon signed-rank test against chance level (0.5, Bonferroni-corrected to account for the number of subjects with an alpha level of 0.05). A non-parametric test was used because it is better suited to handle the non-parametric nature of accuracy values. Then, feature patterns were derived by applying the Haufe transformation to the feature weights (*47*). The Haufe transformation makes feature weights interpretable by converting them into feature patterns that directly reflect the contributions of features’ spatial sources to the model’s predictions. While raw feature weights often mix statistical relevance with noise and are influenced by the properties of the data representation, the transformation projects these weights back to the data space, disentangling the predictive components from the data structure. A feature pattern of zero indicates no contribution from the corresponding neural source, while positive patterns indicate that increased activity in those sources enhances the predicted outcome. Conversely, negative patterns suggest that increased activity in those sources suppresses or inversely relates to the predicted outcome, providing interpretable results. In other words, this process provided, for each behavioral subject and category, interpretable feature importance within the STM space, highlighting the specific regions of the modulation space used to decode each category. For each category, feature patterns were then averaged across behavioral subjects (Figure 1C).

### Identification of STM-sensitive sEEG contacts

#### Ridge regressions

The goal of the analysis was to identify sEEG contacts that exhibited significant responses to the spectrotemporal modulations of the audio track. To achieve this, a ridge regression model was implemented, using the brain signals recorded from each sEEG contact (n=846)— decomposed into six frequency bands (as described above)—as features, and the time course of each spectrotemporal combination of the STM-decomposed audio track (n=2180) as the target variable. To enhance computational efficiency, both the feature and target variables were temporally downsampled to 50 Hz. To account for potential delays between brain responses and audio features, the ridge regression model was applied using a temporal response function (TRF), incorporating time lags ranging from 0 to 1.5 seconds. A backward- modeling approach was used, mapping brain signals onto audio features, as opposed to the more conventional forward-modeling approach. Data were shuffled prior to splitting into folds to ensure robust generalization, and the model was evaluated using 10-fold cross-validation. The optimal regularization parameter (α) was selected by testing values sampled logarithmically across 20 points, spanning from 10^-10^ to 10^10^. Model performance for each fold was assessed using Spearman’s correlation between the predicted (ŷ) and actual (y) time courses of the spectrotemporal combinations. Spearman’s rank correlation was used as a measure of model accuracy because it provides a robust, monotonic estimate of the relationship between predicted and observed STM time courses, without assuming linearity or being overly sensitive to outliers. The final accuracy was calculated as the average correlation across all folds.

For subsequent analyses involving model weight extraction, only the models trained with the optimal regularization parameter, determined based on the highest average accuracy across folds, were used. The extracted weights were then averaged across all time lags tested with the TRF, providing a summary measure of feature importance that integrated responses over the entire lag period.

#### Accuracy map and thresholding

The analysis described earlier resulted in, for each sEEG contact, a correlation coefficient (rho) from a ridge regression model predicting the time course of STM amplitude based on brain signal for each spectrotemporal combination. In other words, an accuracy map of rhos in the STM space was obtained for each sEEG contact (see Figure 2B for an example). After applying a Fisher z-transformation to the rhos, statistical significance was evaluated using one-tailed t-tests for correlation coefficients on each rho of each accuracy map. Correcting for the number of combinations in the STM space, sEEG contacts and folds gives approximately 1.8 × 10⁷ comparisons and a Bonferroni-corrected threshold of approximately 2.7 × 10⁻⁹. We adopted instead a fixed alpha level of 10⁻³⁰, which is more conservative, in order to retain only those contacts for which STM tracking was robust, following a similar procedure in (*92*). Only sEEG contacts showing at least six adjacent significant rhos within the 2D STM space were retained.

### Spatial clustering

To identify brain areas responsive to STM, the MNI coordinates of STM-sensitive sEEG contacts were extracted. For each significant sEEG contact, a 5-mm radius 3D sphere was projected onto an MNI-standardized MRI volume using the MarsBar toolbox (https://marsbar-toolbox.github.io/). The radius was chosen based on the estimated spatial resolution of the bipolar montage (*88*). Only clusters consisting of at least two overlapping sEEG contacts from at least three different patients (to ensure generalizability) were considered significant, while isolated contacts were excluded to minimize Type I error rate.

### Quantifying frequency bands contributions to temporal and spectral feature tracking

To characterize the contribution of each frequency band to the tracking of STM features in STM-sensitive electrodes, we employed a dimensionality reduction approach focused on spectrotemporal combinations relevant to speech (temporal features) and music (spectral features). First, we generated binary masks based on pattern amplitudes from the initial logistic regression models (see *STM features used in perceptual categorization* Methods section). For the temporal dimension, masks included combinations falling within the 90th percentile of pattern amplitudes in speech categorization models; for the spectral dimension, we selected the 90th percentile from music categorization models. We then extracted feature weights from the ridge regression models corresponding to the spectrotemporal combinations identified by these binary masks. For each STM-sensitive electrode and oscillatory band, weights were z-scored across bands, then averaged across the selected spectrotemporal combinations and TRF time lags. This yielded a metric of relative contribution for each frequency band within the temporal and spectral dimensions.

Finally, to assess the relative contribution of frequency bands across dimensions, we conducted a 6×2 non-parametric factorial ANOVA using the Aligned Rank Transform (ART) method (ARTool package in R, *93*). The model included dimension (temporal; spectral) and frequency band (delta; theta; alpha; beta; low gamma; high gamma) as within-subject factors.

Post-hoc pairwise comparisons were performed using Wilcoxon signed-rank tests with False Discovery Rate (FDR) correction for multiple comparisons.

### Using brain-reconstructed STMs to decode behavioral categorization

In this analysis, we extracted from STM-sensitive sEEG contacts the prediction outputs from the previously described ridge regression model. For each sEEG contact, a predicted time-course of the fluctuation of amplitude of each spectrotemporal combination was generated from the corresponding ridge regression model, thus obtaining a predicted audio track in the STM space (i.e., brain-reconstructed STMs). Then, we used the brain-reconstructed STMs as features in a logistic regression model and the binary behavioral judgment of the category (speech/music) as the target variable. The data were shuffled before splitting into folds and the model was evaluated using stratified 5-fold cross-validation. For each category, only sEEG contacts showing significant model accuracy across behavioral subjects were retained, as determined by a Wilcoxon signed-rank test against chance level (0.5) for each category and sEEG contact across behavioral subjects (Bonferroni-corrected to account for the number of sEEG contacts with an alpha level of 0.05). Feature patterns were then derived from significant models by applying the Haufe transformation to the feature weights (*47*) . This resulted in, for each sEEG contact, behavioral subject and category, interpretable feature importance within the STM space. To precisely identify the specific regions of the STM space used to decode each category, one-tailed t-tests (right-tailed) against 0 on z-scored feature patterns were conducted for each combination within the STM space and each sEEG contact across behavioral subjects (Bonferroni-corrected adjusting for the number of sEEG contacts, and combinations, with an alpha level of 0.05). The one-tailed approach was used because only positive feature patterns indicate that increased activity in the feature space enhances prediction outcomes (*47*). This analysis resulted in a representation that indicated, for each point in the STM space, the number of sEEG contacts exhibiting a significant feature pattern (Figure 3E).

### Similarity between feature patterns using brain-reconstructed vs. actual audio track for behavioral categorization

For this analysis, we used two sets of feature patterns. First, we used the feature patterns from the previous logistic regression, where the STMs of all audio track samples served as the features, and the binary behavioral judgment of the category acted as the target variable for each category and participant. Second, we used the feature patterns from the logistic regression where the brain-reconstructed audio track in the STM space served as the features, and the behavioral categorization served as the target variable for each STM-sensitive sEEG contact (for which model accuracy was significantly greater than chance, as described in the previous section), category, and behavioral participant. For the second set, feature patterns from STM-sensitive sEEG contacts were averaged for each hemisphere of the brain. We used spearman correlations to quantify similarity between feature patterns, and their significance was tested using Bonferroni correction to account for the number of participants, categories, and hemispheres, with an alpha level of 0.05. This process resulted in a correlation coefficient for each behavioral participant, category, and hemisphere. In addition to the main analysis described above, we conducted a control analysis to assess the statistical significance of the observed similarity patterns. Specifically, for the first model, in which a logistic regression was applied to binary behavioral judgments using the STM representation of the actual audio track, we disrupted the relationship between predictors and data by shuffling the binary judgments independently for each model. We then performed the same similarity analysis on these shuffled datasets, allowing us to obtain a random distribution of correlation coefficients. To evaluate whether the observed correlation coefficients were significantly different from those obtained with shuffled predictors, we conducted a Wilcoxon signed-rank test for each category, comparing the true distribution to the shuffled distribution of correlation coefficients. Finally, to assess hemisphere preference for each category, we conducted a 2x2 non-parametric factorial ANOVA using the aligned rank transform (ART) method as implemented in the ARTool package in R (*93*) on correlation coefficients, which served as similarity scores. The design included category (speech/music) as a within-subject factor and hemisphere (left/right) as a between-subject factor. Post-hoc comparisons were performed using pairwise Wilcoxon signed-rank tests, with FDR correction applied to control for multiple comparisons.

### Complementary analysis on supplementary dataset

#### Stimuli

The stimulus used in this complementary dataset consisted of a controlled continuous audio track in which speech and music segments were evenly interleaved, in contrast to the more naturalistic Harry Potter audio material, where speech was embedded in conversational and acoustically variable contexts and music spanned for longer consecutive time points than speech (see example Figure 1A). To achieve a clearer contrast between categories, the speech material was selected to provide continuous, uninterrupted monologue, whereas the music was chosen for its sustained and richly structured harmonic content. Specifically, the speech stimulus consisted of a 6 min 49 s excerpt from the French podcast Du jour au lendemain (“Le jour où Anne-Fleur a arrêté "La France”, Mathilde Abassi), and the music stimulus consisted of a 6 min 50 s excerpt from Jehan Alain’s organ solo piece Aria (1938), performed by Jean-Baptiste Robin. These selections were intended to maximize the contrast between speech and music representations in the spectrotemporal modulation space while avoiding any familiarity.

Both audio excerpts were segmented into shorter clips with durations randomly sampled from a truncated normal distribution. For each segment, a target duration was drawn from a Gaussian distribution (μ = 32 s, σ = 4 s), with values constrained within a predefined range (24-40 s) to avoid excessively short or long segments. The continuous audio stream was then cut according to these sampled durations. Fade-ins and fade-outs were applied at the start and end of each segment to avoid clicks. This approach introduced controlled variability in clip length while preserving a consistent temporal scale across stimuli. All segmented excerpts were RMS-normalized to ensure a consistent overall sound level across stimuli.

The 13 excerpts from each category were then manually arranged using REAPER (Cockos Inc.). Segments were placed in strict alternation (i.e., one music excerpt followed by one speech excerpt), while ensuring that each excerpt contained a portion presented in isolation, without overlap from the other category (see Figure S3). This procedure resulted in a balanced temporal structure in which speech-only, music-only, and overlapping segments each accounted for 33% of the total duration. There were no periods of silence. The final composite audio track had a total duration of 10 min 14 s and was compressed using Reaper’s FAST attack compressor with default parameters.

#### Patients

sEEG data from five patients, suffering from drug-resistant focal epilepsy were collected (self-reported biological sex, all females, mean age = 31.8 years, SD = 11.1 years). These patients were distinct from those included in the primary analyses. Participants reported normal hearing. All participants provided written informed consent, and the experimental procedures were approved by the Ethics Review Board of the CIUSSS de la Capitale Nationale (2022– 2476). The study has been conducted according to the principles expressed in the Declaration of Helsinki.

#### Procedure

The five patients underwent the same experimental protocol as the main cohort (see methods section *Procedure: sEEG experiment*).

No behavioral speech/music categorization data were collected, as the temporal structure of the stimuli was fully controlled. Instead, binary time series indicating the presence or absence of speech and music were directly derived from the stimulus design (see Figure S3A).

### Extraction of spectrotemporal modulations

Spectrotemporal modulations were extracted following the same procedure as in the main analysis (see Methods, *Extraction of Spectrotemporal Modulations*), with the only difference that the controlled stimulus was sampled at 44.1 kHz, compared with 48 kHz for the *Harry*

*Potter* audio. This sampling rate yielded 20 temporal modulation points and 100 spectral modulation points, resulting in 2,000 spectrotemporal combinations, slightly fewer than the 2,180 combinations obtained for the *Harry Potter* stimulus.

### sEEG recording and preprocessing

sEEG contact localization, recording, preprocessing, and Hilbert transforms were performed exactly as described for the main analysis (see Methods, *sEEG recording and preprocessing*).

## Supporting information

This PDF includes: Supplementary text, Tables S1 to S3, Figures S1 to S3

## Author contributions

JG: Conceptualization, Data curation, Formal analysis, Investigation, Methodology, Software, Validation, Visualization, Writing – original draft, Writing – review & editing. CG: Data curation, Formal analysis, Software. ECD: Data curation, Formal analysis, Software; AB: Data curation, Software; BM: Writing – review & editing. LM, PLB: Data curation; RJZ: Conceptualization, Funding acquisition, Project administration, Resources, Investigation, Methodology, Supervision, Writing – review & editing, PA: Conceptualization, Data curation, Funding acquisition, Project administration, Resources, Formal analysis, Investigation, Methodology, Validation, Supervision, Writing – review & editing. The manuscript’s contents have been approved by all the co-authors, and they have provided consent for its publication.

## Data, Code, and Materials Availability

All data and code needed to evaluate and reproduce the results in the paper are present in the paper and/or the Supplementary Materials. This study did not generate new materials.

Data used in the figures and the supplementary materials, including auditory stimuli, deidentified and preprocessed sEEG data, and behavioral data, are posted on Open Science Framework (OSF) at https://doi.org/10.17605/OSF.IO/4726S.

We thank the patients for their participation, and Romain Quentin and Oscar Bedford for their valuable discussions and insights.

## REFERENCES

1. H. B. Barlow, Possible principles underlying the transformation of sensory messages. Sens. Commun. 1, 217–233 (1961).

2. W. W. Gaver, What in the World Do We Hear?: An Ecological Approach to Auditory Event Perception. Ecol. Psychol. 5, 1–29 (1993).

3. M. Riesenhuber, T. Poggio, Hierarchical models of object recognition in cortex. Nat. Neurosci. 2, 1019–1025 (1999).

4. J. K. Bizley, Y. E. Cohen, The what, where and how of auditory-object perception. Nat. Rev. Neurosci. 14, 693–707 (2013).

5. T. Chi, P. Ru, S. A. Shamma, Multiresolution spectrotemporal analysis of complex sounds. J. Acoust. Soc. Am. 118, 887–906 (2005).

6. J. P. Rauschecker, S. K. Scott, Maps and streams in the auditory cortex: nonhuman primates illuminate human speech processing. Nat. Neurosci. 12, 718–724 (2009).

7. P. Albouy, L. Benjamin, B. Morillon, R. J. Zatorre, Distinct sensitivity to spectrotemporal modulation supports brain asymmetry for speech and melody. Science 367, 1043–1047 (2020).

8. R. Massoudi, M. M. Van Wanrooij, H. Versnel, A. J. Van Opstal, Spectrotemporal response properties of core auditory cortex neurons in awake monkey. PLoS One 10, e0116118 (2015).

9. B. N. Pasley, S. V. David, N. Mesgarani, A. Flinker, S. A. Shamma, N. E. Crone, R. T. Knight, E. F. Chang, Reconstructing Speech from Human Auditory Cortex. PLoS Biol. 10, e1001251 (2012).

10. F. A. Rodríguez, H. L. Read, M. A. Escabí, Spectral and temporal modulation tradeoff in the inferior colliculus. J. Neurophysiol. 103, 887–903 (2010).

11. R. Santoro, M. Moerel, F. De Martino, R. Goebel, K. Ugurbil, E. Yacoub, E. Formisano, Encoding of Natural Sounds at Multiple Spectral and Temporal Resolutions in the Human Auditory Cortex. PLoS Comput. Biol. 10, e1003412 (2014).

12. M. Schönwiesner, R. J. Zatorre, Spectro-temporal modulation transfer function of single voxels in the human auditory cortex measured with high-resolution fMRI. Proc. Natl. Acad. Sci. 106, 14611–14616 (2009).

13. S. Shamma, On the role of space and time in auditory processing. Trends Cogn. Sci. 5, 340–348 (2001).

14. N. C. Singh, F. E. Theunissen, Modulation spectra of natural sounds and ethological theories of auditory processing. J. Acoust. Soc. Am. 114, 3394–3411 (2003).

15. F. E. Theunissen, K. Sen, A. J. Doupe, Spectral-Temporal Receptive Fields of Nonlinear Auditory Neurons Obtained Using Natural Sounds. J. Neurosci. 20, 2315– 2331 (2000).

16. S. M. N. Woolley, T. E. Fremouw, A. Hsu, F. E. Theunissen, Tuning for spectro- temporal modulations as a mechanism for auditory discrimination of natural sounds. Nat. Neurosci. 8, 1371–1379 (2005).

17. P. Albouy, S. A. Mehr, R. S. Hoyer, J. Ginzburg, Y. Du, R. J. Zatorre, Spectro- temporal acoustical markers differentiate speech from song across cultures. Nat. Commun. 15, 4835 (2024).

18. T. M. Elliott, L. S. Hamilton, F. E. Theunissen, Acoustic structure of the five perceptual dimensions of timbre in orchestral instrument tones. J. Acoust. Soc. Am. 133, 389–404 (2013).

19. A. J. E. Kell, D. L. K. Yamins, E. N. Shook, S. V. Norman-Haignere, J. H. McDermott, A Task-Optimized Neural Network Replicates Human Auditory Behavior, Predicts Brain Responses, and Reveals a Cortical Processing Hierarchy. Neuron 98, 630–644.e16 (2018).

20. J. W. Lewis, W. J. Talkington, A. Puce, L. R. Engel, C. Frum, Cortical Networks Representing Object Categories and High-level Attributes of Familiar Real-world Action Sounds. J. Cogn. Neurosci. 23, 2079–2101 (2011).

21. S. Norman-Haignere, N. G. Kanwisher, J. H. McDermott, Distinct Cortical Pathways for Music and Speech Revealed by Hypothesis-Free Voxel Decomposition. Neuron 88, 1281–1296 (2015).

22. S. Norman-Haignere, J. Feather, D. Boebinger, P. Brunner, A. Ritaccio, J. H. McDermott, G. Schalk, N. Kanwisher, A neural population selective for song in human auditory cortex. Curr. Biol. 32, 1470–1484.e12 (2022).

23. L. Xing, E. Formisano, L. Riecke, Deep Sound Synthesis Reveals Novel Category-Defining Sound Features in the Human Auditory Cortex. BioRxiv [Preprint] (2025). 10.1101/2025.02.14.638230.

24. J. Rissman, A. D. Wagner, Distributed Representations in Memory: Insights from Functional Brain Imaging. Annu. Rev. Psychol. 63, 101–128 (2012).

25. N. Staeren, H. Renvall, F. De Martino, R. Goebel, E. Formisano, Sound Categories Are Represented as Distributed Patterns in the Human Auditory Cortex. Curr. Biol. 19, 498–502 (2009).

26. X. Chen, J. Affourtit, R. Ryskin, T. I. Regev, S. Norman-Haignere, O. Jouravlev, S. Malik-Moraleda, H. Kean, R. Varley, E. Fedorenko, The human language system, including its inferior frontal component in “Broca’s area,” does not support music perception. Cereb. Cortex 33, 7904–7929 (2023).

27. A. D. Friederici, Hierarchy processing in human neurobiology: how specific is it? Philos. Trans. R. Soc. B Biol. Sci. 375, 20180391 (2020).

28. A. Boemio, S. Fromm, A. Braun, D. Poeppel, Hierarchical and asymmetric temporal sensitivity in human auditory cortices. Nat. Neurosci. 8, 389–395 (2005).

29. F. Haiduk, R. J. Zatorre, L. Benjamin, B. Morillon, P. Albouy, Spectrotemporal cues and attention jointly modulate fMRI network topology for sentence and melody perception. Sci. Rep. 14, 5501 (2024).

30. H. L. Jamison, K. E. Watkins, D. V. M. Bishop, P. M. Matthews, Hemispheric Specialization for Processing Auditory Nonspeech Stimuli. Cereb. Cortex 16, 1266– 1275 (2006).

31. P. Robert, R. J. Zatorre, A. Gupta, J. Sein, J.-L. Anton, P. Belin, E. Thoret, B. Morillon, Auditory hemispheric asymmetry for actions and objects. Cereb. Cortex 34, bhae292 (2024).

32. M. Schönwiesner, R. Rübsamen, D. Y. Von Cramon, Hemispheric asymmetry for spectral and temporal processing in the human antero-lateral auditory belt cortex. Eur. J. Neurosci. 22, 1521–1528 (2005).

33. R. J. Zatorre, P. Belin, Spectral and temporal processing in human auditory cortex. Cereb. Cortex 11, 946–953 (2001).

34. J. H. McDermott, E. P. Simoncelli, Sound Texture Perception via Statistics of the Auditory Periphery: Evidence from Sound Synthesis. Neuron 71, 926–940 (2011).

35. D. Boebinger, S. V. Norman-Haignere, J. H. McDermott, N. Kanwisher, Music-selective neural populations arise without musical training. J. Neurophysiol. 125, 2237–2263 (2021).

36. N. J. Zuk, E. S. Teoh, E. C. Lalor, EEG-based classification of natural sounds reveals specialized responses to speech and music. NeuroImage 210, 116558 (2020).

37. D. A. Depireux, J. Z. Simon, D. J. Klein, S. A. Shamma, Spectro-Temporal Response Field Characterization With Dynamic Ripples in Ferret Primary Auditory Cortex. J. Neurophysiol. 85, 1220–1234 (2001).

38. P. W. Hullett, L. S. Hamilton, N. Mesgarani, C. E. Schreiner, E. F. Chang, Human Superior Temporal Gyrus Organization of Spectrotemporal Modulation Tuning Derived from Speech Stimuli. J. Neurosci. 36, 2014–2026 (2016).

39. J. H. Venezia, S. M. Thurman, V. M. Richards, G. Hickok, Hierarchy of speech-driven spectrotemporal receptive fields in human auditory cortex. NeuroImage 186, 647– 666 (2019).

40. N. Sankaran, M. K. Leonard, F. Theunissen, E. F. Chang, Encoding of melody in the human auditory cortex. Sci. Adv. 10, eadk0010 (2024).

41. L. Bellier, A. Llorens, D. Marciano, A. Gunduz, G. Schalk, P. Brunner, R. T. Knight, Music can be reconstructed from human auditory cortex activity using nonlinear decoding models. PLOS Biol. 21, e3002176 (2023).

42. A.-L. Giraud, D. Poeppel, Cortical oscillations and speech processing: emerging computational principles and operations. Nat. Neurosci. 15, 511–517 (2012).

43. J. Giroud, A. Trébuchon, D. Schön, P. Marquis, C. Liegeois-Chauvel, D. Poeppel, B. Morillon, Asymmetric sampling in human auditory cortex reveals spectral processing hierarchy. PLOS Biol. 18, e3000207 (2020).

44. N. Te Rietmolen, M. R. Mercier, A. Trébuchon, B. Morillon, D. Schön, Speech and music recruit frequency-specific distributed and overlapping cortical networks. eLife 13, RP94509 (2024).

45. A. Flinker, W. K. Doyle, A. D. Mehta, O. Devinsky, D. Poeppel, Spectrotemporal modulation provides a unifying framework for auditory cortical asymmetries. *Nat*. Hum. Behav. 3, 393–405 (2019).

46. T. M. Elliott, F. E. Theunissen, The Modulation Transfer Function for Speech Intelligibility. PLoS Comput. Biol. 5, e1000302 (2009).

47. S. Haufe, F. Meinecke, K. Görgen, S. Dähne, J.-D. Haynes, B. Blankertz, F. Bießmann, On the interpretation of weight vectors of linear models in multivariate neuroimaging. NeuroImage 87, 96–110 (2014).

48. A. Borderie, A. Caclin, J.-P. Lachaux, M. Perrone-Bertollotti, R. S. Hoyer, P. Kahane, H. Catenoix, B. Tillmann, P. Albouy, Cross-frequency coupling in cortico-hippocampal networks supports the maintenance of sequential auditory information in short-term memory. PLOS Biol. 22, e3002512 (2024).

49. M. F. Glasser, T. S. Coalson, E. C. Robinson, C. D. Hacker, J. Harwell, E. Yacoub, K. Ugurbil, J. Andersson, C. F. Beckmann, M. Jenkinson, S. M. Smith, D. C. Van Essen, A multi-modal parcellation of human cerebral cortex. Nature 536, 171–178 (2016).

50. N. Ding, A. D. Patel, L. Chen, H. Butler, C. Luo, D. Poeppel, Temporal modulations in speech and music. Neurosci. Biobehav. Rev. 81, 181–187 (2017).

51. J. Giroud, J. P. Lerousseau, F. Pellegrino, B. Morillon, The channel capacity of multilevel linguistic features constrains speech comprehension. Cognition 232, 105345 (2023).

52. J. Giroud, A. Trébuchon, M. Mercier, M. H. Davis, B. Morillon, The human auditory cortex concurrently tracks syllabic and phonemic timescales via acoustic spectral flux. Sci. Adv. 10, eado8915 (2024).

53. A. Chang, X. Teng, M. F. Assaneo, D. Poeppel, The human auditory system uses amplitude modulation to distinguish music from speech. PLOS Biol. 22, e3002631 (2024).

54. J. J. Eggermont, Temporal modulation transfer functions in cat primary auditory cortex: separating stimulus effects from neural mechanisms. J. Neurophysiol. 87, 305–321 (2002).

55. M. A. Escabí, L. M. Miller, H. L. Read, C. E. Schreiner, Naturalistic auditory contrast improves spectrotemporal coding in the cat inferior colliculus. J. Neurosci. 23, 11489–11504 (2003).

56. A. Hsu, S. M. N. Woolley, T. E. Fremouw, F. E. Theunissen, Modulation Power and Phase Spectrum of Natural Sounds Enhance Neural Encoding Performed by Single Auditory Neurons. J. Neurosci. 24, 9201–9211 (2004).

57. L. M. Miller, M. A. Escabí, H. L. Read, C. E. Schreiner, Spectrotemporal Receptive Fields in the Lemniscal Auditory Thalamus and Cortex. J. Neurophysiol. 87, 516–527 (2002).

58. J. H. Venezia, V. M. Richards, G. Hickok, Speech-Driven Spectrotemporal Receptive Fields Beyond the Auditory Cortex. Hear. Res. 408, 108307 (2021).

59. D. R. Langers, W. H. Backes, P. V. Dijk, Spectrotemporal features of the auditory cortex: the activation in response to dynamic ripples. NeuroImage 20, 265–275 (2003).

60. D. Scheinost, S. Noble, C. Horien, A. S. Greene, E. Mr. Lake, M. Salehi, S. Gao, X. Shen, D. O’Connor, D. S. Barron, S. W. Yip, M. D. Rosenberg, R. T. Constable, Ten simple rules for predictive modeling of individual differences in neuroimaging. NeuroImage 193, 35–45 (2019).

61. P. Albouy, J. Mattout, R. Bouet, E. Maby, G. Sanchez, P.-E. Aguera, S. Daligault, C. Delpuech, O. Bertrand, A. Caclin, B. Tillmann, Impaired pitch perception and memory in congenital amusia: the deficit starts in the auditory cortex. Brain 136, 1639–1661 (2013).

62. P. Loui, D. Alsop, G. Schlaug, Tone Deafness: A New Disconnection Syndrome? J. Neurosci. 29, 10215–10220 (2009).

63. B. Milner, Laterality effects in audition. Interhemispheric Relat. Cereb. Domin., 177– 195 (1962).

64. H. Okamoto, R. Kakigi, Hemispheric Asymmetry of Auditory Mismatch Negativity Elicited by Spectral and Temporal Deviants: A Magnetoencephalographic Study. Brain Topogr. 28, 471–478 (2015).

65. A. J. Sihvonen, T. Särkämö, A. Rodríguez-Fornells, P. Ripollés, T. F. Münte, S. Soinila, Neural architectures of music – Insights from acquired amusia. Neurosci. Biobehav. Rev. 107, 104–114 (2019).

66. 66. R. J. Zatorre, From Perception to Pleasure: The Neuroscience of Music and Why We Love It (Oxford University PressNew York, ed. 1, 2024; https://academic.oup.com/book/55154).

67. X. Teng, X. Tian, J. Rowland, D. Poeppel, Concurrent temporal channels for auditory processing: Oscillatory neural entrainment reveals segregation of function at different scales. PLOS Biol. 15, e2000812 (2017).

68. K. Cha, R. J. Zatorre, M. Schönwiesner, Frequency Selectivity of Voxel-by-Voxel Functional Connectivity in Human Auditory Cortex. Cereb. Cortex 26, 211–224 (2016).

69. E. B. J. Coffey, S. C. Herholz, A. M. P. Chepesiuk, S. Baillet, R. J. Zatorre, Cortical contributions to the auditory frequency-following response revealed by MEG. Nat. Commun. 7, 11070 (2016).

70. K. L. Hyde, I. Peretz, R. J. Zatorre, Evidence for the role of the right auditory cortex in fine pitch resolution. Neuropsychologia 46, 632–639 (2008).

71. R. J. Zatorre, Hemispheric asymmetries for music and speech: Spectrotemporal modulations and top-down influences. Front. Neurosci. 16, 1075511 (2022).

72. C. E. Shannon, A mathematical theory of communication. Bell Syst. Tech. J. 27, 379–423 (1948).

73. D. J. Graham, D. J. Field, Efficient neural coding of natural images. New Encycl Neurosci 1, 1–18 (2007).

74. E. P. Simoncelli, B. A. Olshausen, Natural Image Statistics and Neural Representation. Annu. Rev. Neurosci. 24, 1193–1216 (2001).

75. J. Gervain, M. N. Geffen, Efficient Neural Coding in Auditory and Speech Perception. Trends Neurosci. 42, 56–65 (2019).

76. M. S. Lewicki, Efficient coding of natural sounds. Nat. Neurosci. 5, 356–363 (2002).

77. V. L. Ming, L. L. Holt, Efficient coding in human auditory perception. J. Acoust. Soc. Am. 126, 1312–1320 (2009).

78. E. C. Smith, M. S. Lewicki, Efficient auditory coding. Nature 439, 978–982 (2006).

79. R. D. Patterson, S. Uppenkamp, I. S. Johnsrude, T. D. Griffiths, The Processing of Temporal Pitch and Melody Information in Auditory Cortex. Neuron 36, 767–776 (2002).

80. M. A. Goodale, A. D. Milner, Separate visual pathways for perception and action. Trends Neurosci. 15, 20–25 (1992).

81. R. Vogels, G. A. Orban, “Chapter 14 Coding of stimulus invariances by inferior temporal neurons” in Progress in Brain Research (Elsevier, 1996; https://linkinghub.elsevier.com/retrieve/pii/S0079612308633300)vol. 112, pp. 195–211.

82. A. Feinsinger, N. Pouratian, H. Ebadi, R. Adolphs, R. Andersen, M. S. Beauchamp, E. F. Chang, N. E. Crone, J. L. Collinger, I. Fried, A. Mamelak, M. Richardson, U. Rutishauser, S. A. Sheth, N. Suthana, N. Tandon, D. Yoshor, Ethical commitments, principles, and practices guiding intracranial neuroscientific research in humans. Neuron 110, 188–194 (2022).

83. J. Peirce, J. R. Gray, S. Simpson, M. MacAskill, R. Höchenberger, H. Sogo, E. Kastman, J. K. Lindeløv, PsychoPy2: Experiments in behavior made easy. Behav. Res. Methods 51, 195–203 (2019).

84. J. L. Fleiss, B. Levin, M. C. Paik, Statistical Methods for Rates and Proportions (Wiley, ed. 1, 2003; https://onlinelibrary.wiley.com/doi/book/10.1002/0471445428) *Wiley Series in Probability and Statistics*.

85. M. Gamer, J. Lemon, I. F. P. S. <[email protected]>, Irr: Various Coefficients of Interrater Reliability and Agreement (2019; https://CRAN.R-project.org/package=irr).

86. C.-C. Huang, E. T. Rolls, J. Feng, C.-P. Lin, An extended Human Connectome Project multimodal parcellation atlas of the human cortex and subcortical areas. Brain Struct. Funct. 227, 763–778 (2022).

87. F. Tadel, S. Baillet, J. C. Mosher, D. Pantazis, R. M. Leahy, Brainstorm: A User-Friendly Application for MEG/EEG Analysis. Comput. Intell. Neurosci. 2011, 1–13 (2011).

88. K. Jerbi, T. Ossandón, C. M. Hamamé, S. Senova, S. S. Dalal, J. Jung, L. Minotti, O. Bertrand, A. Berthoz, P. Kahane, J. Lachaux, Task-related gamma-band dynamics from an intracerebral perspective: Review and implications for surface EEG and MEG. Hum. Brain Mapp. 30, 1758–1771 (2009).

89. A. Gramfort, M. Luessi, E. Larson, D. A. Engemann, D. Strohmeier, C. Brodbeck, R. Goj, M. Jas, T. Brooks, L. Parkkonen, M. S. Hämäläinen, MEG and EEG Data Analysis with MNE-Python. Front. Neurosci. 7, 1–13 (2013).

90. F. Pedregosa, G. Varoquaux, A. Gramfort, V. Michel, B. Thirion, O. Grisel, M. Blondel, P. Prettenhofer, R. Weiss, V. Dubourg, J. Vanderplas, A. Passos, D. Cournapeau, M. Brucher, M. Perrot, E. Duchesnay, Scikit-learn: Machine Learning in Python. J. Mach. Learn. Res. 12, 2825–2830 (2011).

91. P. Guilleminot, M. Thornton, E. Varano, M. Kegler, sPyEEG: A Stimuli-Brain Mapping Toolbox for Electrophysiological Data Analysis. (2026).

92. J. Berezutskaya, M. J. Vansteensel, E. J. Aarnoutse, Z. V. Freudenburg, G. Piantoni, M. P. Branco, N. F. Ramsey, Open multimodal iEEG-fMRI dataset from naturalistic stimulation with a short audiovisual film. Sci. Data 9, 91 (2022).

93. L. A. Elkin, M. Kay, J. J. Higgins, J. O. Wobbrock, “An Aligned Rank Transform Procedure for Multifactor Contrast Tests” in The 34th Annual ACM Symposium on User Interface Software and Technology (ACM, Virtual Event USA, 2021; https://dl.acm.org/doi/10.1145/3472749.3474784), pp. 754–768.

