## Supplementary material for "Auditory cortical coding of spectrotemporal modulations is sufficient to support perceptual categorization of speech and music": This PDF includes: Supplementary text, Tables S1 to S3, Figures S1 to S3

**Table S1: Coordinates of STM-sensitive sEEG contacts**. Labels were extracted from the extended Human Connectome Project (HCPex) multimodal parcellation atlas (*86*), a modified and expanded version of the HCP-MMP v1.0 atlas (*49*).

| Hemisphere | Cluster number | MNI coordinates [x y z] | HCPex label | Patient number |
|---|---|---|---|---|
| Left | 1 | [-45 -26 -6] | Area STSd posterior | 1 |
| | | [-53 -24 -1] | Auditory 5 Complex | 1 |
| | | [-62 -22 -1] | Area STSd posterior | 1 |
| | | [-43 -21 -6] | Area STSd posterior | 7 |
| | | [-50 -20 -6] | Area STSd posterior | 7 |
| | | [-57 -19 -7] | Auditory 5 Complex | 7 |
| | | [-63 -18 -4] | Auditory 5 Complex | 7 |
| | | [-66 -17 -2] | Auditory 5 Complex | 7 |
| | | [-65 -17 -6] | Auditory 5 Complex | 11 |
| | 2 | [-50 -7 -4] | Medial Belt Complex | 1 |
| | | [-56 -5 -4] | Auditory 5 Complex | 1 |
| | | [-61 -2 -5] | Auditory 5 Complex | 1 |
| | | [-45 -2 -17] | Para Insular Area | 2 |
| | | [-52 -1 -14] | Area STSd anterior | 2 |
| | | [-57 -1 -12] | Auditory 5 Complex | 2 |
| | | [-47 -10 -10] | Para Insular Area | 3 |
| | | [-53 -7 -10] | Area STSd anterior | 3 |
| | | [-58 -5 -11] | Auditory 5 Complex | 3 |
| | | [-63 -4 -13] | Area TE1 anterior | 3 |
| | | [-44 -3 -12] | Para Insular Area | 7 |
| | | [-57 0 -10] | Auditory 5 Complex | 7 |
| | | [-60 2 -10] | Auditory 5 Complex | 7 |
| Right | 3 | [55 0 -11] | Auditory 5 Complex | 1 |
| | | [60 2 -10] | Auditory 5 Complex | 1 |
| | | [63 3 -9] | Auditory 5 Complex | 1 |
| | | [47 -12 -5] | Para Insular Area | 3 |

| | | | | |
|---|---|---|---|---|
| | | [54 -11 -5] | Area STSd anterior | 3 |
| | | [61 -11 -5] | Auditory 5 Complex | 3 |
| | | [65 -11 -4] | Auditory 5 Complex | 3 |
| | | [53 -1 -10] | Auditory 5 Complex | 5 |
| | | [58 1 -7] | Auditory 5 Complex | 5 |
| | | [64 1 -6] | Auditory 5 Complex | 5 |
| | | [68 -5 -2] | Auditory 4 Complex | 8 |
| | 4 | [36 -19 6] | Area_52 | 3 |
| | | [39 -24 -2] | L Belt | 5 |
| | | [44 -24 2] | L Belt | 5 |
| | | [49 -26 5] | L Belt | 5 |
| | | [55 -28 8] | L Belt | 5 |
| | | [53 -20 3] | Auditory 5 Complex | 6 |

**Figure S1: Spatial distribution of STM-sensitive sEEG contacts** across left, frontal, and right brain views

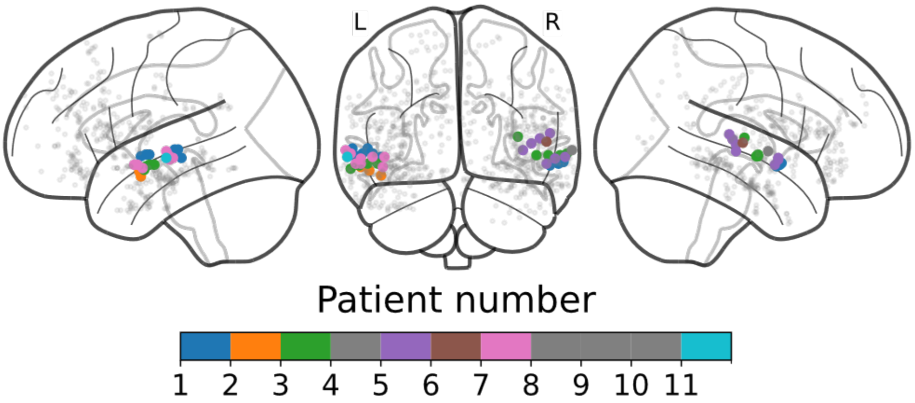


**Table S2: Detailed post-hoc analysis results of the frequency band-by-dimension interaction.** Statistics derive from the 6x2 aligned rank test (ART) on z-scored feature weights with frequency band (delta; theta; alpha; beta; low-gamma; high-gamma) and dimension (temporal; spectral) as within subject factors. FDR-corrected p-values are represented in aqua for the temporal dimension, in orange for the spectral dimension, and in bold when significant ($p$ < 0.05).

| | Delta | Theta | Alpha | Beta | Low Gamma | High Gamma |
|---|---|---|---|---|---|---|
| Delta | | 0.5 | **5.4e-04** | 0.046 | **1.18e-04** | **8.25e-11** |
| Theta | 0.075 | | **1.18e-04** | **1.70e-02** | **5.36e-04** | **1e-10** |
| Alpha | **1.10e-06** | **0.005** | | 0.17 | **2.95e-10** | **7.70e-14** |
| Beta | 0.93 | 0.12 | **1.04e-05** | | **8.72e-08** | **9.68e-13** |
| Low Gamma | **8.85e-06** | **5.32e-08** | **2.46e-12** | **1.04e-05** | | **2.17e-04** |
| High Gamma | **0.03** | **8.32e-04** | **4.24e-07** | **0.021** | 0.17 | |

**Figure S2: Accuracy from logistic regression models predicting behavioral categorization with brain-reconstructed STM features.** For each STM-sensitive electrode, model accuracy is shown for predicting behavioral binary time series (n = 19 participants) using reconstructed STMs as features, separately for the speech (top panel) and music (bottom panel) categories. Asterisks indicate electrodes for which model performance was significantly above chance (chance = 0.5; $p < 0.05$, FDR-corrected), based on Wilcoxon signed-rank tests performed across behavioral vectors (n = 19) for each electrode and category.

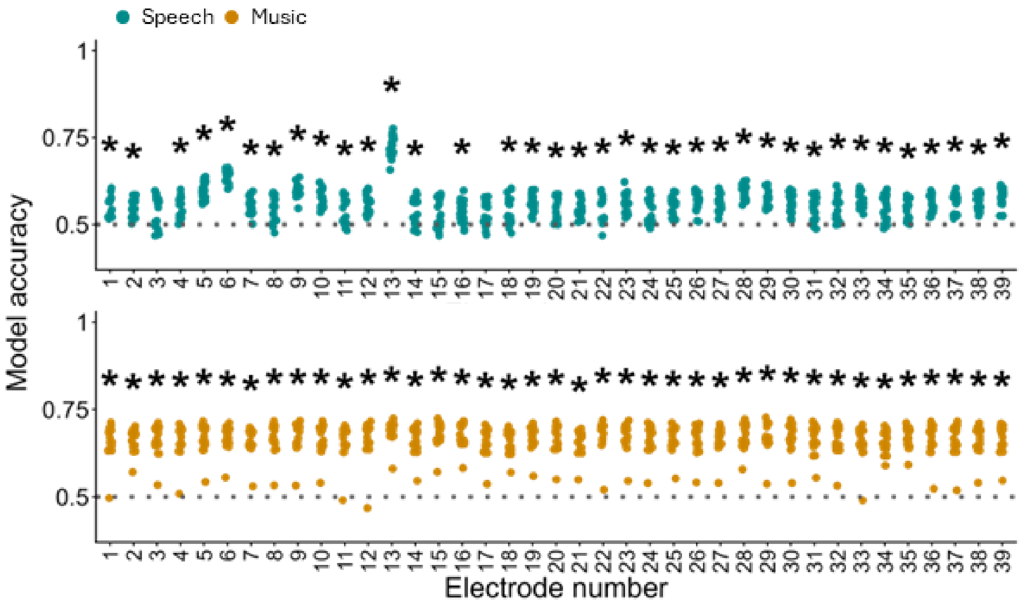


**Detailed results from the complementary analysis**

STM features predict perceptual categorization

We performed the same logistic regression analysis as in the corresponding main result section, with the distinction that only one model per category was fitted. This was possible because no behavioral data were required for the controlled stimulus, since the binary time series indicating the presence or absence of speech and music were directly derived from the stimulus design. Both models achieved 99% accuracy, consistent with the results obtained using the Harry Potter stimulus.

Feature patterns closely resembled those from the main analysis (Figure S3B). For the speech category, the STM features most important for the binary classification mainly contained temporal modulations. The 90th percentile of the highest feature values spanned 2.4-15 Hz in temporal modulations and only 0-1.18 cyc/kHz in spectral modulations (dotted line, Figure S3B, left). Peak activity (99th percentile; solid line) was concentrated at 5.6–8 Hz and 10.4-12.8 Hz temporally, and 0-0.6 cyc/kHz spectrally. For the music category, spectral modulations were the primary distinguishing features. The 90th percentile of high activity covered most of the spectral modulation range (1.36–9 cyc/kHz) and 0–2.4 Hz temporally (dotted line, Figure S3B, right), while peak activity (99th percentile; solid line) was concentrated at 6.8–7.15 cyc/kHz and 7.6–8.06 cyc/kHz spectrally, and 0–0.8 Hz temporally.

<u>Bilateral non-primary auditory areas track spectrotemporal modulations \</u>

This analysis was performed following the exact same procedure as described in the corresponding section of the main results. It should be noted that the five patients who completed the task with the controlled stimulus were distinct from those included in the main analysis and exhibited a different implantation scheme, with very few contacts in the left hemisphere (Figure S3C). After computing accuracy maps from the mTRF ridge models linking STM features to oscillatory power, thresholding these maps, and clustering significant electrodes (see Methods), we identified a single cluster of 23 STM-sensitive contacts in the right non-primary auditory area (see Table S3 for MNI coordinates and atlas labels). No clusters were observed in the left hemisphere, consistent with the sparse left-hemisphere coverage in this patient set. Within the right-hemisphere cluster, contacts were located in regions already identified in the main analysis, primarily the posterior distal STS, areas A5 and A4, and the parabelt. Overall, these results demonstrate that, in a completely independent set of patients and using markedly different speech and music stimuli, sEEG contacts in non-primary auditory regions reliably track spectrotemporal modulation features of complex sounds.

<u>Ground-truth speech and music labels can be decoded from brain-reconstructed STMs</u>

This analysis was performed following the exact same procedure as described in the corresponding section of the main results, with the key difference that ground-truth speech and music labels were directly available from the controlled stimulus. Accordingly, one logistic regression model was fitted per electrode to predict the presence or absence of speech and music based on brain-reconstructed STMs. Because inter-subject variability in behavioral responses was no longer a factor, model performance was assessed descriptively across electrodes for each category (Figure S3E). In addition, Wilcoxon signed-rank tests against chance level (0.5) were performed across electrodes, revealing that logistic regression models significantly predicted both speech and music presence/absence (both $p < 0.001$) solely from brain-reconstructed STMs. Taken together, these results replicate our main finding in an independent dataset using a distinct, fully controlled stimulus.

Brain-reconstructed and acoustic STM representations are highly similar

This analysis was performed following the same procedure as described in the corresponding section of the main results, with the distinction that, for each electrode and each category, similarity was assessed between feature patterns derived from the logistic regression based on brain-reconstructed STMs against only one (vs 19 in the main section) obtained from the original logistic regression using the acoustic STMs. As expected, STM patterns were highly similar (Spearman correlation; $p < 0.001$, Bonferroni-corrected for the number of categories and electrodes). This result was further confirmed at the group level (across electrodes) by comparing the distribution of correlation coefficients to a shuffled distribution (see Methods and Figure S3F), revealing highly significant differences for each category (both $p < 0.001$). Note that the hemispheric comparison performed in the main analysis could not be conducted here, as this patient cohort had insufficient electrode coverage in the left hemisphere.

Overall, these complementary analyses fully replicate the main findings in an independent set of patients and with a different stimulus, thereby further strengthening the robustness of our conclusions.

**Figure S3. Complementary analysis.**

(A) Envelope of the controlled stimulus and corresponding ground-truth binary time series for speech and music. The stimulus consisted of speech and music excerpts (~32 s, truncated Gaussian sampling) arranged in strict alternation, yielding a continuous audio stream with balanced proportions of speech-only, music-only, and overlapping segments (33% each).

(B) Feature patterns from the logistic regression predicting ground-truth binary time series with STM features for speech and music. Threshold-based contours highlight regions of high feature patterns amplitude: the 90th percentile of the distribution is outlined with a dotted line, and the 99th percentile is outlined with a solid line.

(C) Full sEEG coverage of the five patients in standard MNI space on the left and right sides of a glass half-brain visualization.

(D) STM-sensitive contacts. Peak $R^2$ values from contacts that survived statistical thresholding and spatial clustering (see Methods). One cluster (n = 23 contacts) was identified in the right non-primary auditory cortex (see Table S3 for coordinates).

(E) Accuracy from logistic regression models predicting behavioral categorization with brain-reconstructed STM features for each STM-sensitive electrode and each category.

(F) Feature pattern similarity distribution. Density curves represent the distribution of correlation coefficients between feature patterns derived from the two models depicted in Figure 4A. The top (aqua) distribution corresponds to the speech category, and the bottom (orange) distribution corresponds to the music category. For comparison, a separate distribution (gray) shows the correlation coefficients obtained when feature patterns from the first model (top panel of Figure 4A) were compared with those from the second model (bottom panel of Figure 4) after shuffling the behavioral categorization labels.

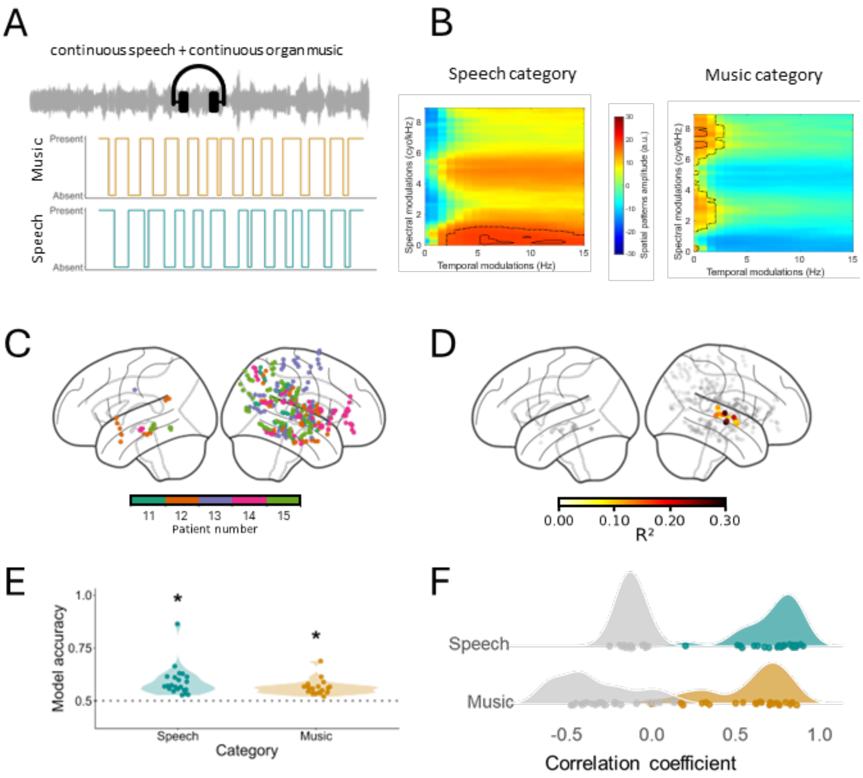


**Table S3: Coordinates of STM-sensitive sEEG contacts for the complementary analysis on the controlled audio track.** Labels were extracted from the extended Human Connectome Project (HCPex) multimodal parcellation atlas (*86*), a modified and expanded version of the HCP-MMP v1.0 atlas (*49*).

| Hemisphere | Cluster number | MNI coordinates [x y z] | HCPex label | Patient number |
|---|---|---|---|---|
| Right | 1 | [53 -14 1] | Area STSd posterior | 12 |
| | | [50 -16 -4] | Area STSd posterior | 13 |
| | | [57 -17 -5] | Auditory 5 Complex | 13 |
| | | [64 -16 -3] | Auditory 5 Complex | 13 |
| | | [49 -4 -7] | Area TA2 | 14 |
| | | [54 -5 -3] | Area TA2 | 14 |
| | | [ 61 -6 -2] | Auditory 4 Complex | 14 |
| | | [67 -8 1] | Auditory 4 Complex | 14 |
| | | [37 -23 3] | Area 52 | 14 |
| | | [50 -24 4] | Area STSd posterior | 14 |
| | | [56 -25 5] | Area STSd posterior | 14 |
| | | [63 -28 3] | Auditory 5 Complex | 14 |

| | | | | |
|---|---|---|---|---|
| | | [44 -21 4] | Auditory 5 Complex | 15 |
| | | [51 -20 4] | Auditory 5 Complex | 15 |
| | | [57 -19 4] | Auditory 5 Complex | 15 |
| | | [64 -18 5] | Auditory 5 Complex | 15 |
| | | [68 -18 6] | Auditory 5 Complex | 15 |
| | | [54 -5 -6] | Area STSd anterior | 16 |
| | | [61 -6 -5] | Auditory 5 Complex | 16 |
| | | [64 -6 -5] | Auditory 5 Complex | 16 |
| | | [55 -26 10] | Parabelt Complex | 16 |
| | | [62 -26 10] | Parabelt Complex | 16 |
| | | [66 -26 12] | Auditory 4 Complex | 16 |
